# Methylglyoxal is the main culprit to impairing neuronal function: mediated through tryptophan depletion

**DOI:** 10.1101/2023.03.29.534483

**Authors:** Md. Samsuzzaman, Jae Hyuk Lee, Seong-Min Hong, Hyun jun Park, Keun-A Chang, Hyun-Bum Kim, Myoung Gyu Park, Hyeyoon Eo, Myung Sook Oh, Sun Yeou Kim

## Abstract

Depression is a common and prevalent illness and the exact cause of major depressive disorder is not known. Here, we investigated how methylglyoxal (MGO) stress induces depression and unveiled the potential molecular mechanism. Our *in vivo* results suggested that MGO caused depression in mice, confirmed by several behavioral tests. Interestingly, it halted the mice’s brain’s tryptophan levels and its related neurotransmitters. In addition, MGO induced a reduction in the number of cells in different hippocampal regions. Moreover, it decreased tryptophan hydroxylase 1 (TPH1) and tryptophan hydroxylase 1 (TPH2) levels in the brain and large intestine. Surprisingly, MGO showed the highest affinity and trapping ability toward tryptophan. Most importantly, combined treatment with MGO-tryptophan displayed similar effects as those exhibited by the tryptophan-null treatment in neuronal cells, which included neuronal apoptosis, decrease TPH1 and TPH2 levels, and inhibition of neuronal outgrowth. However, tryptophan treatment improved MGO induced depression-like behavior of mice and recovered the loss of neuronal and hippocampal cells. Subsequently, it also induced MGO detoxifying factors, tryptophan levels, and reduces inflammation in the intestine. Collectively, our data revealed that MGO induced depression facilitated by neuronal and synaptic dysfunction is mediated through the disturbance of tryptophan metabolism in the brain and intestine.

## Introduction

Diabetes mellitus (DM) is the most common and recognizable disease that causes mortality and morbidity in South Asian and Western communities [1]. Depression is another root cause of mortality and disease complications [2]. Studies on depressive disorder patients and animals indicate the possible underlying factors for this condition, such as impaired neuronal connectivity, decreased neuron spine density, and hippocampal cell loss [3, 4, 5, 6]. The bidirectional relations between DM and depression have been demonstrated by many research groups. Based on Meta-analysis, the depressive disorder has been significantly and consistently associated with many types of diabetic complications. Patients with DM are at a high risk of depressive disorder, and in addition, also have a greater chance of developing DM [7, 8]. Moreover, depression is highly prevalent in patients with diabetic complications [9]. One statistical analysis found a correlation between depression and poorly controlled hyperglycemic conditions [10]. Importantly, the effect of long-term hyperglycemia and diabetic complications in depression should be checked. Nevertheless, DM and depressive disorder are closely associated with chronic hyperglycemia, hyperinsulinemia, insulin resistance, and hypoglycemic conditions in diabetic patients [11]. Hyperglycemia causes cognitive decline, neurodegeneration, dementia, and brain aging. The pathogenic mechanisms of brain damage due to hyperglycemia are too complex and involve a combination of vascular disease, oxidative stress, mitochondrial dysfunction, apoptosis, neuroinflammation, decrease in neurotrophic factors, changes in neurotransmitter level, and accumulation of amyloid β and tau phosphorylation [12, 13, 14]. Interestingly, depressive disorder is more common in patients with Type-2 Diabetes than in those with Type 1 Diabetics [15]. To date, many studies have been done in this regard, but the results have been inconsistent [2, 16, 17, 18, 19]. However, previous studies have provided an ideal basis for the correlation between hyperglycemia and depressive disorders, patients with depression are often addicted to high-carbohydrate diets. In this context, the actual role of hyperglycemia in brain disease is a critical open issue.

In chronic hyperglycemia, highly reactive metabolites produced during glycolysis are the main cause of impaired normal cell function, resulting in different types of chronic diseases [20]. Among them, methylglyoxal (MGO) is the most reactive dicarbonyl substance, it usually binds to the amino acids, especially, arginine and lysine, to form advanced glycation end-products (CEL, MGH1, etc) which ultimately deteriorate normal protein functions [21]. MGO has been reported to play a critical role in different central nervous system (CNS) cognitive functions, particularly anxiety, stress, depression, and neurodegenerative diseases [22]. Moreover, studies have found a high presence of MGO-derived advanced glycation end-products in the brain fluid of a patient with Alzheimer’s disease [23]. Additionally, it has been demonstrated that the extrinsic administration of MGO to cells or rats induces behavioral or physiological changes [24]. Thus, clinical investigations suggest that high levels of MGO in the serum are closely associated with rapid cognitive decline in elderly patients [25]. Despite these interesting shreds of evidence, the actual mechanism of MGO-induced depressive disorder is not yet clear.

Tryptophan is an essential amino acid that needs to be availed by animals from diets through feeding and, slightly in part, from internal protein disruption. The original function of tryptophan is in protein synthesis, which participates in protein-membrane building [26, 27]. If there is a decrease in tryptophan levels, it will impair its targeted protein production, ultimately pulling back cell growth [27]. One of the most important roles of tryptophan is to act as a precursor for neurotransmitters such as serotonin (5-hydroxytryptamine), which has a strong role in mood, stress response, appetite, and especially in depression [26]. The effects of tryptophan depletion on depression and behavioral changes have been investigated in clinical and preclinical studies [28]. Clinical studies revealed that patients with lower levels of tryptophan showed depressive and mood-changing characteristics, although the role of tryptophan in depression remains controversial reported by research groups [29, 30, 31, 32]. Moreover, tryptophan depletion elevated anxiety, immobility, and mood change parameters, as demonstrated in an animal model. The gut-brain axis is now a topic of research interest that represents the relationship between the brain and gut, including the autonomic and enteric nervous systems, as well as the gastrointestinal microenvironment. However, any disturbance in gut microbiota networks in the intestine may lead to severe diseases such as inflammatory bowel syndrome (IBS), which could further cause colorectal cancer [33]. In the past decades, mounting evidence has suggested that microbiota markedly affects the generation and functions of metabolites, which may impact the contribution of the CNS to brain mood and behavior development [34]. Studies have found that tryptophan is a common agent used by the gut microbiota in the production of serotonin (5-HT) and melatonin, in addition to important neurotransmitters such as dopamine (DA), epinephrine (EP), and norepinephrine (NE) [35, 36, 37]. Nevertheless, MGO is also a major bacterial product during the anaerobic glycolysis of carbohydrates in the large intestine. Yet, the actual mechanism of MGO from the intestine in depression and behavioral changes is largely unknown.

In this study, we aimed to understand whether MGO can induce depression and intestinal disorder in mice or not, and, we are looking to find out whether that neuronal dysfunction by MGO treatment is mediated through tryptophan depletion or not. Therefore, we hypothesized that MGO may interrupt tryptophan availability, which triggers neuronal cell death; thus, affecting depressive behaviors in mice, apart from having a great impact on the intestine.

## Materials and Methods

### Materials

Material and common methods information details are provided in the Additional file 1.

### Animals aend study design

ICR mice (7-weeks-old, male) were acquired from Orient Bio (Gyeonggi-do, Korea) and placed in a standard temperature-(22±2°C) with 65% humidity and a 12 h/12 h light/dark cycle. The Animal Care Committee of the Center of Animal Care and Use at Gachon University (GIACUC-R2020008) reviewed and approved all experimental guidelines. After the one week of the acclimation period, the ICR mice were indiscriminately divided into four groups: control group (C, n=8), MGO 25 mg/kg (n=8), MGO 30 mg/kg (n=8), and MGO 65 mg/kg (n=8). MGO was administered to the experimental ICR mice for 2 weeks at different doses (25, 30, and 65 mg/kg) via rectal injection (30% v/v glycerol in phosphate buffer saline).

### Measurement of tryptophan, 5-HTP, and 5-HT levels in the plasma

Tryptophan, 5-HTP, and 5-HT levels were analyzed according to Takada et al with slight modifications [40]. Briefly, mouse plasma was added to 0.1% formic acid and acetonitrile (CAN), vortexed, and mixed within 30 s for 5 min. The reaction samples were then centrifuged, vortexed with 0.1% formic acid, and injected into the Agilent LC 1100 series LC-MS/MS system (CA, USA). Chromatography was performed using an Agilent ZORBAX Extend-C18 column (1.0 × 150 mm, 3.5 micron) operated at 30°C. The solvent system consisted of mobile phase A [100% acetonitrile (ACN)] and mobile phase B (0.1% formic acid in distilled water) at a ratio of 1:1 (v/v). To detect tryptophan, 5-HTP, and 5-HT in the analytes, the MS/MS system was operated under positive (ESI+), negative (ESI-), and multiple reaction monitoring modes.

### Measurement of neurotransmitter levels in brain tissues

The levels of DA, NE, EP, and 5-HT were analyzed using the protocol reported by Bishnoi et al [41]. After the sacrifice of mice, the whole brain was removed and homogenized in 0.1 M perchloric acid (PCA) (10 mg/μL). The samples were then spin down at 12,000 rpm for 30 min. The supernatant was filtered using 0.2 μm filters and injected into the HPLC system (Waters Corp., Massachusetts (MA), USA). The samples (20 μL) were analyzed using a Kromasil C18 column (150 mm × 4.6 mm, 5 µm) with an electrochemical detector. A mobile phase consisting of 2% citric acid, 2% K_2_HPO_4_, 1 mM EDTA, 1.2% MeOH, and 7 mg/mL sodium octyl sulfate was used. The detector conditions were +0.008 V and a sensitivity range of 0-100 nA. The flow rate was maintained at 1.0 mL/min.

### Measurement of MGO and GO levels in mixtures and plasma

MGO and Glyoxal (GO) levels were analyzed following the reported protocol with slight modifications [42]. The standard of unreacted MGO and GO in the samples was analyzed in terms of the peak area of quinoxaline and 2-methyl quinoxaline (2-MQ) to that of 5-methyl quinoxaline (5-MQ). It has been reported that o-phenylenediamine (*o*-PD) reacts with GO to form quinoxaline. The plasma samples were incubated with 0.45 N PCA and 10 mM *o*-PD for 24 h at room temperature. The reaction samples were then centrifuged, filtered, and injected into an HPLC system. The samples were analyzed using a Kromasil C18 column (250 mm × 4.6 mm, 5 µm) with 20% ACN as the mobile phase, at a detection wavelength of 315 nm. The flow rate was maintained at 1.0 mL/min.

### Histological analysis and immunohistochemistry

Paraffin blocks of the large intestine, small intestine, and brain tissues were fixed in 10% formalin. After fixation, all tissues were dehydrated by exposure to ethanol and washed with xylene, and embedded in paraffin blocks following sectioned to 5 µm thickness. The sections were stained using a hematoxylin and eosin (H&E) staining kit (Sigma-Aldrich). For immunohistochemical analysis, the sections were incubated in 0.1% protease K in phosphate buffer saline (PBS) for antigen retrieval and incubated in 3% H_2_O_2_ in PBS for 15 min. The sections were then incubated with 1% normal horse serum in PBS for 20 min. After blocking, the sections were incubated with TPH1 and TPH2 overnight at 4°C in a shaker. The sections were washed and incubated with biotinylated respective immunoglobulin G (IgG) secondary antibodies at room temperature (25°C) for 1 h. The sections were then rinsed with PBS and incubated with VECTASTAIN^®^ ABC reagent for 30 min. Antibody expression was detected with the help of 3,3-diaminobenzidine (DAB). Nuclei were counterstained with Hoechst 33342 for 5 min. The stained tissue slides were observed and photographed under a Nikon Eclipse 80i microscope (Nikon, Tokyo, Japan) at 100× magnification.

### Multi-electrode array recording

The 8 x 8 microelectrode array (MEA) has two temperature-control units that keep the solution and baseplate at 33 °C within 0.01 °C, an amplifier, and a quadruple channel stimulus generator. The MEA was brushed, coated with 0.1 percent polyethylenimine (PEI; Sigma), and UV sterilized for at least three hours after being soaked in 2 percent ultrasonol 7 (Carl Roth GmbH, Karlsruhe, Germany) for one hour. Probes were cleaned with 2 percent ultrasonol 7 in distilled water for 30 minutes in between studies, then rinsed and maintained in distilled water at room temperature. Rat brain slices were presoaked in artificial cerebrospinal fluid (CSF), which was centered on the MEA and perfused at a rate of 3 ml/min. The artificial CSF included 124 NaCl, 26 NaHCO_3_, 10 glucose, 3 KCl, 2 CaCl_2_, 1 MgCl_2_, and 10 HEPES (in mM). It was also adjusted to pH 7.4. Slices were stabilized at 33 °C for one hour. Slice and MEA array was mounted on a MEA1060 amplifier interface (1,200 dB gain), and the interface was grounded using an Ag/AgCl pellet. A desktop computer handled 60 channels of data sampled at 25 kHz using MC Rack and MC Data Tool (Multi Channel Systems, GmbH).

### Cell culture and viability assay

BV2, N2a, and SH-SY5Y cells were cultured in high-glucose Dulbecco’s modified Eagle medium supplemented with 10% fetal bovine serum and 1% penicillin/streptomycin and maintained at 37°C in a humidified incubator containing 5% CO_2_. Human intestinal epithelial cells (HIEC-6) were cultured in Opti-MEM™ I Reduced Serum Medium (Thermo Scientific, MA, USA). Cell viability was measured using MTT assays (ref). Cells were seeded and treated with different concentrations (10, 100, 250, 500, 750, and 1000 µM) of MGO for an additional 24 h. The absorbance was measured at 570 nm using a microplate reader (Molecular Devices, CA, USA). The morphology images were obtained using the IncuCyte^®^ ZOOM imaging system (Essen BioScience, Michigan (MI), USA).

### Cell migration assay

Cells were seeded into 96-well plates for 24 h, following which the confluent monolayers of the cultured cells were scratched with a Woundmaker tool (Essen BioScience). The culture medium was removed, and cells were washed with PBS before being treated with different concentrations (250, 500, 750, and 1000 µM) of MGO for 24 h. The scratched cells were observed every 2 h using the IncuCyte^®^ ZOOM Imaging System.

### Cell apoptosis assay

Apoptosis and necrosis cell quantity was measured using the FITC/Annexin V Apoptosis Detection Assay Kit I (BD Biosciences Pharmingen, San Diego, CA, USA), according to the manufacturer’s protocol. Briefly, N2a cells were seeded into a 6-well plate (4.0×10^5^ cells/well) and incubated for 24 h. After incubation, the cells were treated with different concentrations (250 and 500 µM) of MGO and tryptophan-free medium for 24 h. The cells were then washed with cold PBS and re-suspended in 1× binding buffer. Following that, 5 µL FITC Annexin V and propidium iodide were added and maintained on a dark clean bench at room temperature (25°C) for 15 min. The treated samples were examined using flow cytometry (FACSCalibur™; Becton, Dickinson and Company, San Jose, CA, USA) within 1 h.

### Neurite outgrowth assay

N2a cells were seeded into a 6-well plate (4.0×10^5^ cells/well) and incubated for 4 h. After incubation, the cells were treated with different concentrations (250 and 500 µM) of MGO and tryptophan-free medium for 24 or 48 h. IncuCyte^®^ ZOOM Imaging System was used to evaluate the neurite outgrowth, neurite length, and cell morphology of treated or untreated cells every 2 h.

### MGO-affinity assay

The MGO-affinity assay was performed to analyze the reaction at different stages of the MGO-amino acid compound process, according to the method of Ina Nemet et al with slight modifications [43]. Briefly, MGO in the presence or absence of amino acids were incubated in PBS (pH 7.4) and 0.02% sodium azide for 7 d at 37°C in the dark. The affinity of MGO-amino acid was evaluated in terms of fluorescence at excitation/emission wavelengths of 355/460 nm, as detected using a VICTOR™ X3 multilabel plate reader (PerkinElmer, MA, USA).

### MGO-trapping assay

The MGO-trapping ability was analyzed using the MGO-trapping assay according to Navarro et al with slight modifications [44]. Briefly, the MGO solution (1.0 mM) was dissolved in PBS (pH 7.4) and incubated with or without tryptophan (1.0 mM) for 1 d and 7 d at 37°C in the dark. After incubation, 10 mM *o*-PD was incubated with the treated or untreated samples for 24 h at room temperature (25°C) in the dark. The reaction samples were then centrifuged, filtered, and injected into an HPLC system. The samples were analyzed using a Kromasil C18 column (250 mm × 4.6 mm, 5 µm) with 20% ACN as the mobile phase at a detection wavelength of 315 nm. The flow rate was 1.0 mL/min and the column temperature was maintained at 30°C.

### Measurement of tryptophan levels in cell culture media and extract

Tryptophan levels were analyzed according to a previously reported method [45]. N2a cells were seeded in a 60 φ dish, washed with cold PBS, centrifuged, and extracted with ice-cold 50% methanol/50% piperazine –N (PIPES)-EDTA at -20°C for 5 to 10 min. After the cell extracts were separated using centrifugation, the samples were injected into an HPLC system. The samples were analyzed using a Sepax HP-C18 column (250 mm × 4.6 mm, 5 µm) with 15 mM potassium phosphate and 2.7% ACN as an isocratic mobile phase at a detection wavelength of 220 nm. The flow rate was 0.8 mL/min and the column temperature was maintained at 30°C.

### Immunofluorescence analysis

N2a cells were seeded into 2-well plates using a glass slide and incubated for 24 h at 37°C in 5% CO_2_. After incubation for 24 h, the cells were treated with different concentrations (500 and 750 µM) of MGO and tryptophan-free medium for 24 h. The cells were then washed with PBS, fixed in 10% formalin for 10 min, and permeabilized with 0.1% Triton™ X-100 for 10 min. After that, the primary antibody was added to each well and the cells were incubated with it for 24 h at 4°C in a shaker. After incubation for 24 h, the cells were washed with PBS, incubated with rabbit anti-goat IgG H&L (Alexa Fluor^®^ 555-conjugated) secondary antibody for 1 h and Hoechst 33342 for 5 min, and then mounted with Fluoromount™ aqueous mounting medium (Sigma-Aldrich). The stained cells were observed under a laser scanning confocal microscope (Nikon A1+) and analyzed using NIS-Elements imaging software.

### Fluorescence immunostaining and Spine density analysis

Hippocampal neurons were fixed in a 4% PFA solution for 20 min at room temperature and washed using 1x PBS. Fixed hippocampal neurons were permeabilized in PBS containing 0.1% Triton X-100 for 15 min and blocked 1% BSA for 45 min at room temperature. Hippocampal neurons were stained with primary antibodies for 1hr at room temperature followed by fluorescence-conjugated secondary antibodies or Phalloidin 488 (Thermo) for 2 hr at room temperature. The hippocampal neurons were washed with PBS and mounted on the sliding glass. Images of hippocampal neurons were captured by a Zeiss LSM 700 confocal microscope with a 20x objective. Dendritic spines were analyzed with Zen 2.3 SP1 software.

### Quantitative real-time PCR

Total RNA was extracted from cultured N2a cells using TRIzol^®^ reagent (Invitrogen, CA, USA). A total of 1 µg of RNA was reverse transcribed to complementary DNA using the PrimeScript™ RT reagent Kit (Takara Bio, Otsu, Japan), according to the manufacturer’s protocol. Quantitative polymerase chain reaction (qPCR) was performed using SYBR^®^ Premix Ex Taq™ (Takara Bio) on an Mx3005P qPCR System (Agilent Technologies). Primer sequences are listed in the supplementary material (Table. S1). Glyceraldehyde 3-phosphate dehydrogenase (*GAPDH)* was used as a housekeeping gene.

### Microarray analysis

N2a cells were seeded into 6-well plates and incubated for 24 h. After incubation for 24 h, the cells were treated with different concentrations (250 and 500 µM) of MGO and tryptophan-free medium for 24 h and extracted with TRIzol^®^ reagent. The prepared total RNA was then sent to Macrogen (Macrogen, Seoul, Korea) for microarray analysis. Microarray analysis was performed using an Agilent Mouse genome microarray (Agilent Technologies, CA, USA), according to the manufacturer’s protocol.

### Statistical analysis

All data have been represented as mean ± standard error of the mean. All statistical analyses were conducted by using Prism 5.0 (GraphPad Software Inc., CA, USA). Statistical comparisons between the control and experimental groups were performed using Bonferroni’s test for multiple comparisons of one-way analysis of variance. Turkey postdoc comparison tests after One-way ANOVA was used to calculate the statistical significance of cell viability assay and spine density analysis. Differences were considered statistically significant at *p*<0.05.

## Results

### MGO treatment enhanced depression/anxiety-like behavior in ICR mice

To clarify whether MGO can affect the sensory and motor functions of ICR mice, we performed several anxiety and depression-like behavior tests. Results of open filed test (OFT), the most extensively used method to check anxiety behavior, revealed that MGO (25, 30, 65 mg/kg)-treated groups displayed a significant reduction in the time spent in the central zone, as compared to the control group (Fig. 1A, D). Less time spent by the mice in the central area indicated fearful characteristics, as it has already been established that depressive subjects do not spend more time in the central zone [46]. Upon performing tail suspension test (TST), another technique to assess depression-like activity in ICR mice, the results confirmed that MGO (25, 30, 65 mg/kg) induced more immobility time in the mice, as compared to that in the control group (Fig. 1B). Similarly, we also detected that MGO (30 and 65 mg/kg) triggered immobility time in ICR mice in the forced swimming test (FST) (Fig. 1C). Therefore, the behavioral test data suggested the depression-and anxiety-inducing ability of MGO in mice. Adrenergic/nor-adrenergic, dopaminergic, and serotonergic systems have been the central areas of pathological studies of depression and/or mood-linked disorders. In this context, we measured DA, EP, NE, and 5-HT levels in mouse brains. As shown in Fig. 1E-H, MGO decreased the DA and E levels in the mouse brain at 30 and 65 mg/kg, and NE at 25 and 65 mg/kg as compared to that in the control, while the 5-HT levels were decreased only at 65 mg/kg, thus indicating that MGO induced a decrease in the neuronal activity.

**Fig. 1:**
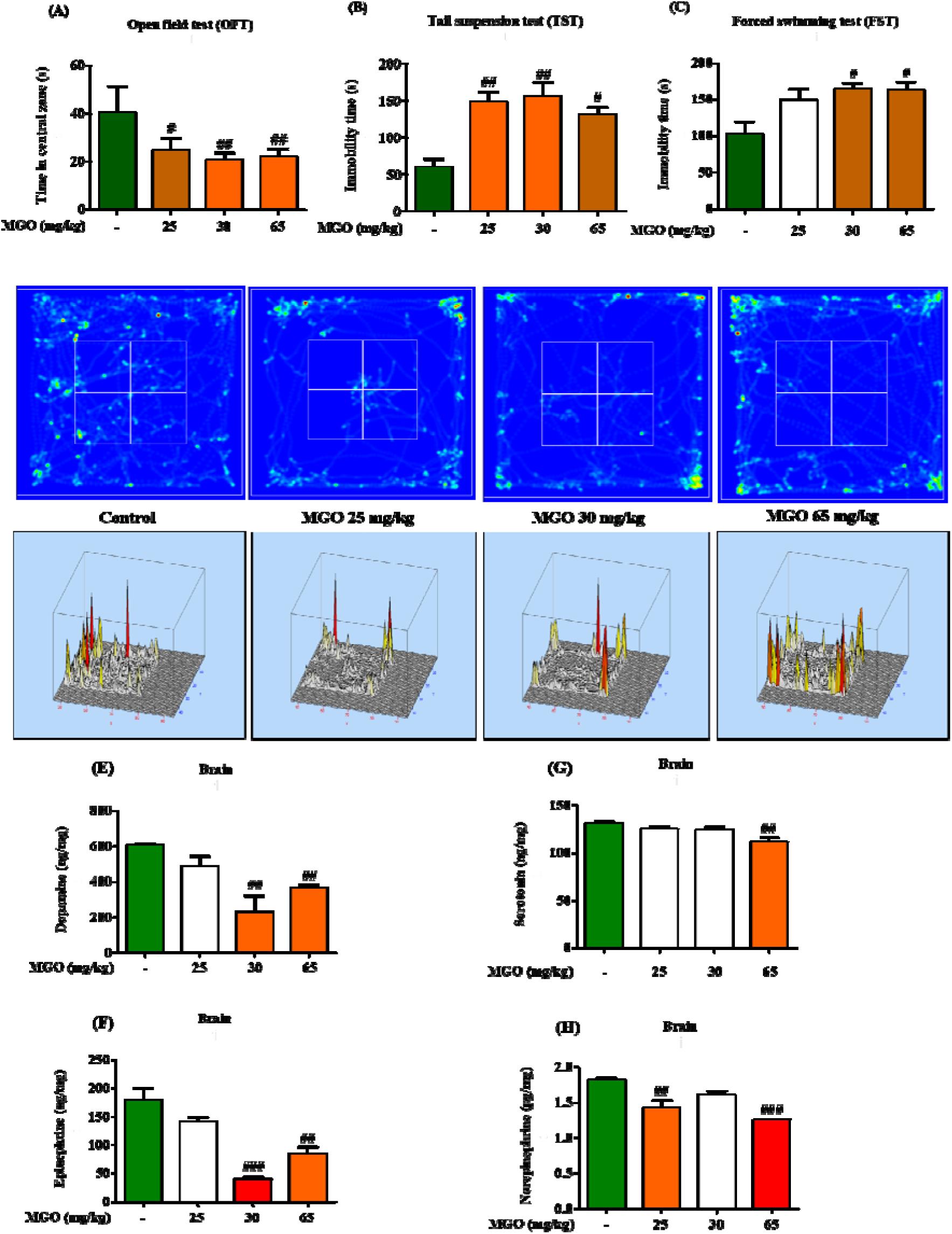
MGO induced depression-like behavior, and reduced tryptophan metabolism-related factors, and neurotransmitters in mice. (A) OFT was done in mice after 3 weeks of MGO administration by the rectal route. Mice were exposed to the open field for 8 min, to examine the time spent in the central zone. (B) Mice were pre-adapted for 2 min before being allowed into the TST chamber for 4 min in the TST. Immobility time was calculated using SMART3.0 SUPER PACK. (C) Mice were pre-adapted for 2 min before being allowed into the FST chamber for 4 min in the FST. Immobility time was calculated using SMART3.0 SUPER PACK. All data have been represented as mean ± SEM. n=5 (^#^*p*<0.05 and ^##^*p*<0.01 *vs.* control). (D) Visual recording of the OFT was carried out using SMART3.0 SUPER PACK. (E-H) Levels of neurotransmitter-related factors (dopamine, epinephrine, norepinephrine, and serotonin) were analyzed using HPLC analysis. The whole brain was removed and homogenized in 0.1 M PCA (10 mg/μL). Following that, the samples were centrifuged at 12,000 rpm for 30 min. The supernatant was filtered using 0.2 μm filters and injected into the HPLC system. All data have been represented as mean ± SEM. n=5 (^##^*p*<0.01 and ^###^*p*<0.001 *vs.* control).

### MGO treatment altered the dentate gyrus (DG), cornu ammonis 3 (CA3), cornu ammonis 1 (CA1) regions, and long-term potentiation (LTP) of the hippocampi of ICR mice

A reduced number of cells and structural changes in the hippocampus have been found to be vitally connected to major depressive disorder [46]. Hence, we postulated that MGO might have negative effects on cell growth in different parts of the hippocampus. Therefore, we investigated the effects of MGO treatment on the DG, CA3, and CA1 regions of the hippocampi of mice. We found that MGO significantly lowered the growth of cells in the DG, CA3, and CA1 regions of the hippocampus at the dose of 65 mg/kg (Fig. 2A-D), as compared to that in the control, which suggests that MGO induced structural changes in the hippocampi of mice. We also investigated the effect of different concentrations of MGO (5 and 100 μM) on hippocampal CA1 LTP. Interestingly, the time-dependent change in field excitatory post-synaptic potential (fEPSP) activity and the mean % of fEPSP from 30 to 40 min after TBS were pooled for analysis. Treatment of MGO at 5 µM increased post-TBS-stimulated fEPSP (169.38±8.03%), as compared to that of the control group (146.96±6.36%). However, a relatively high concentration of MGO (100 µM) inhibited the induction of LTP by TBS (113.14±4.33%) (Fig. 2E-F). Similar to the effects seen upon treatment with 100 µM MGO, induction of LTP was completely blocked upon treatment with cyanquixaline (CNQX) (99.18±4.47%), an AMPA receptor antagonist, with no significant difference compared to the treatment with 100 µM MGO. Therefore, our results suggested that treatment with high-concentration MGO blocks LTP induction in hippocampal tissues.

**Fig. 2:**
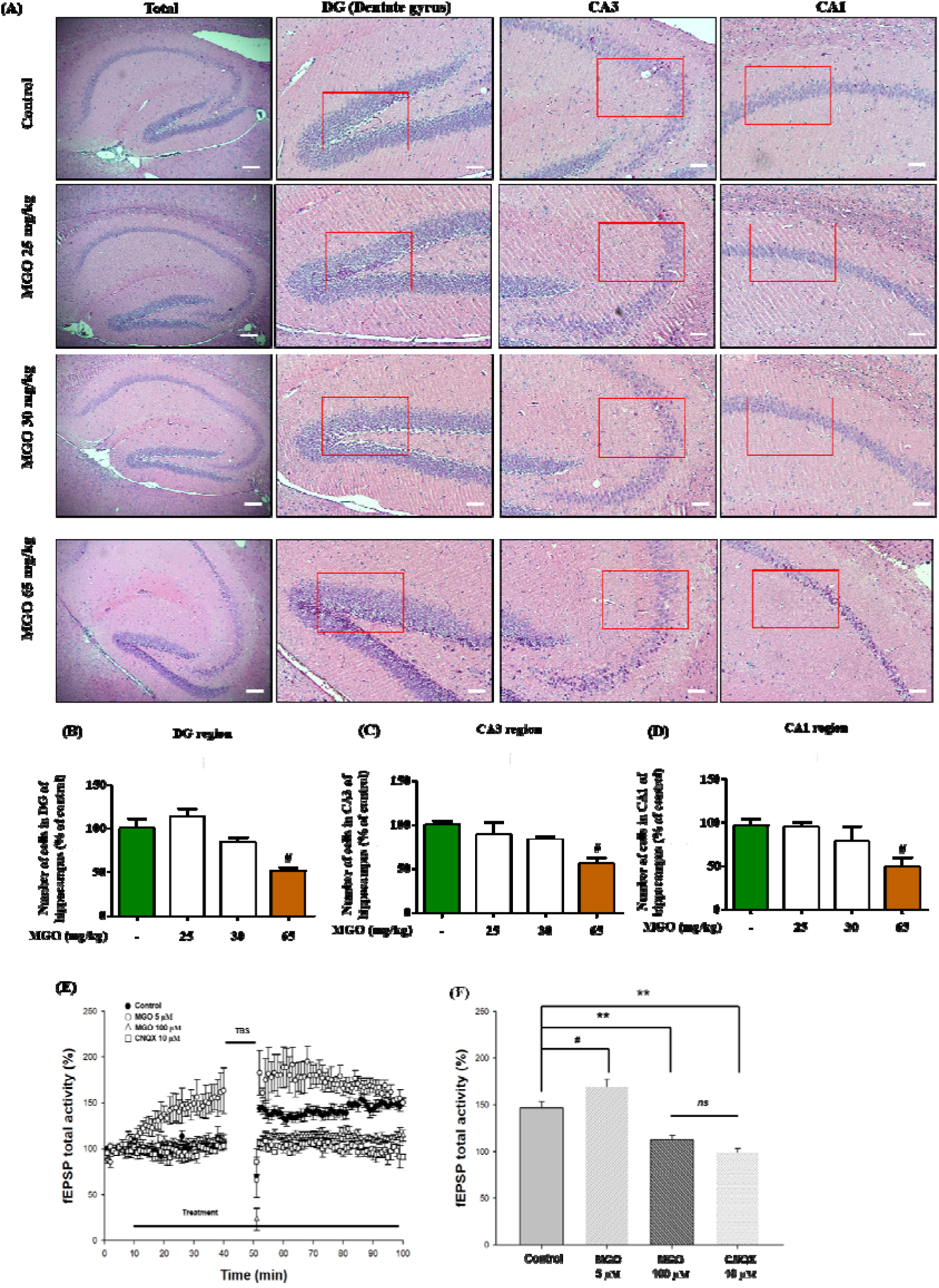
MGO facilitated hippocampal degeneration and lowered the total field excitatory post-synaptic potential (fEPSP) activity. (A) Histological staining of brain sections was carried out. Representative H&E-stained images have been shown in the total hippocampal and dentate gyrus (DG), CA3, and CA1 regions. Scale bar: 100 µm. (B-D) Quantitative measurements of neuronal cells were performed by calculating the number of cells in the DG, CA3, and CA1 regions from selected fields per image. The values were calculated using ImageJ software. All data have been represented as mean ± SEM. n=3 (^#^*p*<0.05 *vs.* control). (E) Long-term potentiation (LTP) from all recordings in the control, MGO-, and CNQX-treated hippocampus. (F) The fEPSP total activity from 30 to 40 min after TBS in the control, MGO-, and CNQX-treated hippocampus. All data have been represented as mean ± S.E.M. Control, treated with nothing after 10 min of the baseline; MGO, treated with 5 and 100 μM of MGO after 10 min of the baseline; CNQX, treated with 10 µM of CNQX after 10 min of the baseline. \*\**p*<0.01 and ^#^*p*<0.1 *vs*. control group using one-way ANOVA with Tukey’s HSD post-hoc test.

### MGO treatment influenced tryptophan-related essential factors in the plasma and hippocampal area of ICR mice

Tryptophan depletion is known to accelerate depression [47], and this notion forced us to hypothesize that MGO-induced depression in mice might be mediated by modulating the levels of tryptophan and its related factors, such as 5-HTP, 5-HT, TPH1, and TPH2, in mice. Thus, we examined the effects of MGO on these parameters using an LC-MS/MS system and immunohistochemistry analysis. Mouse plasma was subjected to LC-MS/MS analysis of tryptophan, 5-HTP, and 5-HT. We found that MGO markedly downregulated tryptophan levels at doses of 30 and 65 mg/kg, and those of 5-HTP and 5-HT in a dose-dependent manner in the mouse plasma (Fig. 3A-C). We then inspected TPH1 and TPH2 expression in different sections of the hippocampus, because lower levels of tryptophan were also found to be associated with impaired hippocampal function. As illustrated in Fig. 3D, while MGO lowered TPH2 expression in the CA1 and CA3 sections at the dose of 65 mg/kg, and in the cortex at doses of 30 and 65 mg/kg, as compared to that in the control group, it did not affect TPH2 expression in the DG section. Fig. 3 E-H represents the statistical analysis of the staining intensity of TPH2 in the DG, CA3, CA1, and cortex sections. We also analyzed TPH1 expression in the cortex section of the mouse brain via immunostaining, as mood may depend on an integral pathway linked to the cortex. Anatomically, the cortex has a direct connection to the hippocampal part of the brain, and our experimental results showed that MGO notably induced the downregulation of TPH1 expression even at the dose of 25 mg/kg in the cortex (Fig. 3I). Fig. 3J represents the statistical analysis of Fig. 3I. These data indicate that the MGO-induced depressive effects in mice might be due to an impaired tryptophan metabolic pathway.

**Fig. 3:**
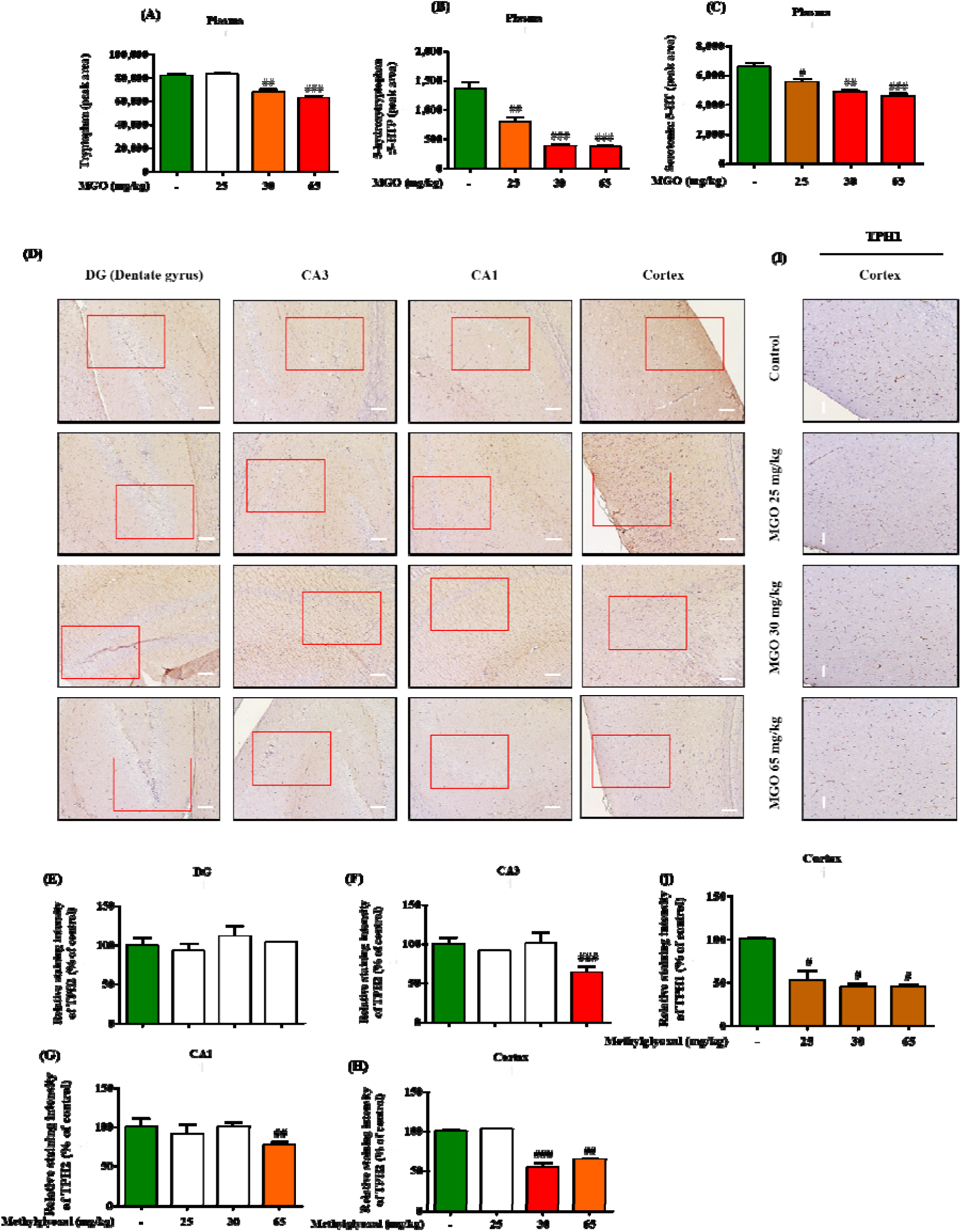
MGO suppressed tryptophan metabolism protein in the plasma, hippocampus, and cortex of mice. (A-C) Levels of tryptophan metabolism-related factors (tryptophan, 5-HTP, and 5-HT) in the mice plasma were analyzed using an LC-MS/MS system. Mice plasma was added to 0.1% formic acid and CAN, vortexed, and mixed within 30 secs for 5 min. Following that, the reaction samples were separated by means of centrifugation, vortexed with 0.1% formic acid, and injected into the LC-MS/MS system. All data have been represented as mean ± SEM. n=3 (^#^*p*<0.05 ^##^*p*<0.01 and ^###^*p*<0.001 *vs.* control). (D) Immunohistochemical (IHC) analysis of brain sections was performed to examine TPH2 levels in the DG, CA3, and CA1 regions. Stained images represent TPH2 levels in the DG, CA3, and CA1 indicated in the box. Scale bar: 100 µm. (E-H) Quantitative measurements of TPH2 intensity were conducted by calculating the expression in the DG, CA3, and CA1 from selected fields per image. The values were calculated using ImageJ software. All data have been represented as mean ± SEM. n=3 (^#^*p*<0.05, ^##^*p*<0.01 and ^###^*p*<0.001 *vs.* control).(I) Immunohistochemical (IHC) analysis of brain sections was performed to examine TPH1 levels in the cortex region. Stained images represent TPH1 levels in the cortex regions indicated in the box. Scale bar: 100 µm. (J) Quantitative measurements of TPH1 intensity were conducted by calculating the expression in the cortex from selected fields per image. The values were calculated using ImageJ software. All data have been represented as mean ± SEM. n=3 (^#^*p*<0.05 *vs.* control).

### MGO treatment regulated tryptophan metabolism and IBS in the colon of ICR mice

Evidence indicates that IBS helps in triggering depressive disorders [48]. Indeed, when we analyzed the colon length and inflammatory cytokines, we found a surprisingly visible decrease in colon length in response to MGO treatment at a dose of 65 mg/kg, as compared to the colon length in the control group, without any effect on the mouse body weight (Fig. 4A and B). Interestingly, we observed that GO levels in the colon were higher upon 65 mg/kg of MGO injection than those in the control group (Fig. 4C); this may be due to the conversion of MGO into GO, which is also a chronic toxic malonaldehyde. It is well established that the gut and brain play an intriguing role in depression and anxiety [49]. As the colon is also one of the major sources of MGO produced by bacteria, we next investigated the role of MGO on TPH1 and TPH2 expression, which are responsible for the conversion of tryptophan to 5-HT, in mouse colon sections. Our immunohistochemical staining experiment revealed that MGO significantly decreased TPH1 expression in the colon, as compared to that in the control colon section (Fig. 4D-F) at doses of 30 and 65 mg/kg, with no effect on TPH2 activity. We also confirmed MGO-induced toxicity and inhibition of wound healing in colon cells (Fig. S1A, B). Moreover, MGO administration significantly increased levels of pro-inflammatory cytokines Tumor necrosis factor-α (TNF-α) and interleukin-6 (IL-6) (Fig. 4H-J), as they are important markers of IBS [50].

**Fig. 4:**
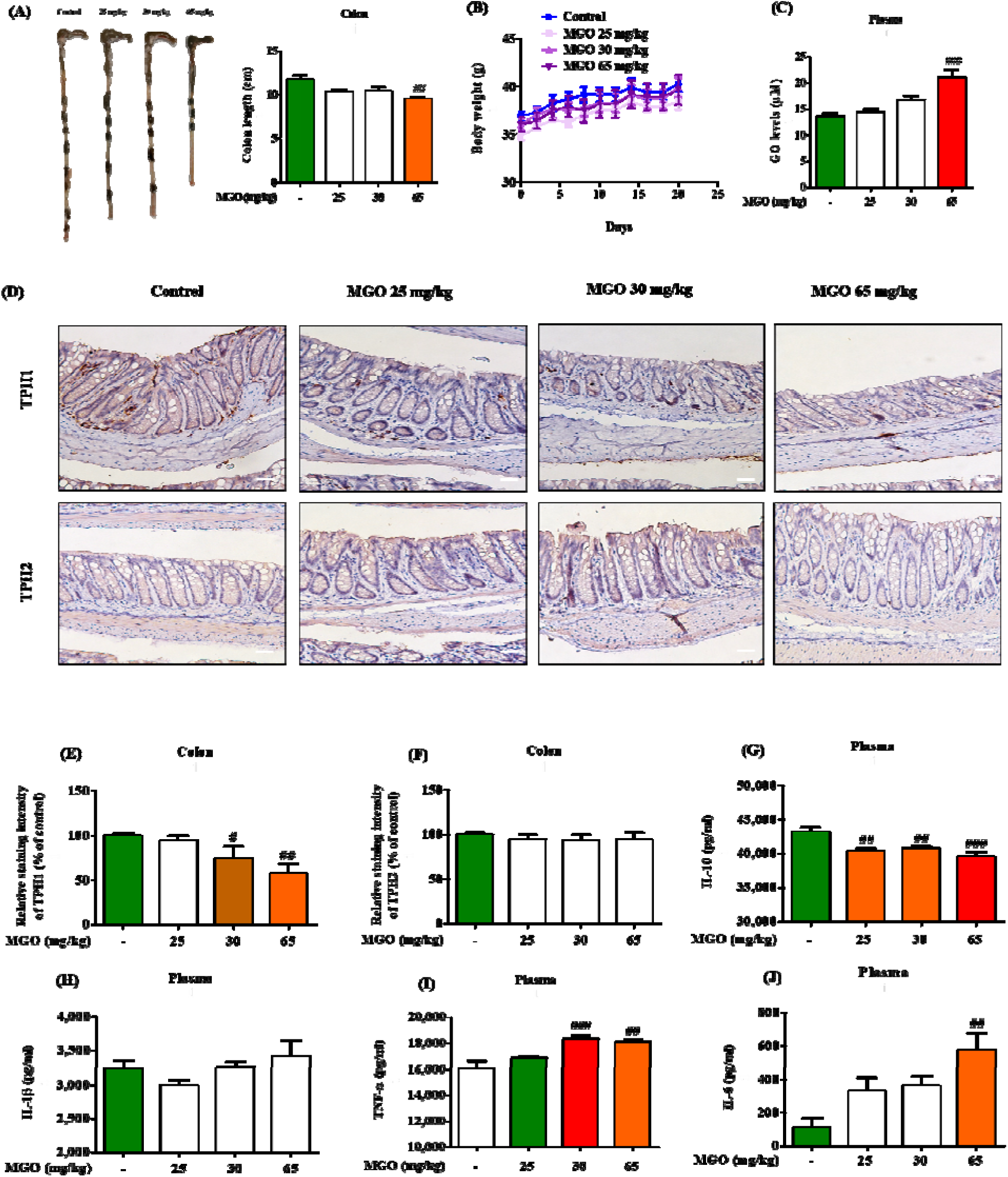
MGO modulated colon length and tryptophan metabolism protein expression in mice colon tissue. (A) The normal control group received only 30% v/v glycerol in PBS, while the three other groups received MGO diluted in PBS. After sacrifice, the colon was measured using a ruler (in cm). Data have been represented as mean ± SEM. n=8 (^##^*p*<0.05 *vs.* control). (B) Bodyweight measurement. (C) The levels of GO in the plasma were analyzed using HPLC. Data have been represented as mean ± SEM. n=8 (^###^*p*<0.001 *vs.* control). (D) Immunohistochemical (IHC) analysis of colon sections was performed. Representative images were stained for TPH1 and TPH2 in the colon tissues. Scale bar: 100 µm. (E, F) Quantitative measurements of TPH1 and TPH2 were conducted by calculating the expression in the colon from selected fields per image. The values were calculated using ImageJ software. Data have been represented as mean ± SEM. n=8 (^#^*p*<0.01^##^*p*<0.001 *vs.* control). (G-J) The secretion of IL-10, TNF-α, IL-1β, and IL-6 cytokines in the plasma was evaluated using the respective ELISA assay kit. All data have been represented as mean ± SEM. n=8 (^##^*p*<0.01 and ^###^*p*<0.001 *vs.* control).

Additionally, MGO significantly decreased the IL-10 levels in the mouse plasma (Fig. 4G). As previously reported, IL-10 is a cytokine with significant anti-inflammatory properties that plays a vital role in the immune response to pathogens, thus maintaining normal tissue homeostasis and preventing damage to the host [51]. Additionally, we checked Zonula occludens-1 (ZO-1) and Ki67 levels in colon tissue to support the MGO induced decreased colon length results. It is known that ZO-1 is one of the most important proteins for intestinal barrier integrity and Ki67, a cell proliferation marker, is an abnormally expressed protein that is associated with bowel inflammation [76]. MGO treatment significantly reduced ZO-1 and increased Ki67 expression levels as compared to control in colon tissue (Fig. S4A, B). These results uncovered another mechanism of MGO-induced depression in mice, that is, by affecting colon homeostasis, including tryptophan and inflammatory pathways.

### MGO showed the highest affinity and trapping-ability towards tryptophan itself, not Arginine/Lysine among other amino acids

Amino acids directly regulate neurotransmitter synthesis, and major depressive disorders are often characterized by decreased levels of amino acids [52]. To determine whether MGO has any correlation with these amino acids, we inspected MGO affinity to all amino acids *via* an affinity assay. Among the tested amino acids, MGO showed the highest affinity for tryptophan, followed by that for histidine, threonine, serine, carnosine, lysine, and arginine (Fig. 5A). The affinity of MGO for tryptophan was approximately seven times that of the normal sample. Based on these data, we observed tryptophan levels in N2a cell culture media and cell extracts in the presence of two concentrations (250 and 500 µM) of MGO. Although tryptophan levels were not affected by MGO in the cell culture media, there was a significant dose-dependent reduction in tryptophan levels only in the cell extract (Fig. 5A and B), as compared to the levels in the control (Table. S2). We used tryptophan-free media as conditioned media. Moreover, tryptophan was chosen for further study because it indicated the most powerful reactive effect against MGO-induced affinity. Therefore, we selected MGO (10 mM) and tryptophan (1.0 mM) sample mixtures for further analysis, to verify our affinity assay data through an MGO-trapping assay using HPLC. As shown in Fig. 5C, dramatic differences were detected in tryptophan concentration between the two groups (1 d and 7 d) of the MGO and tryptophan mixture (Table. S3). On day one, the tryptophan concentration detected was 239.51±13.38 µM, which was further reduced to 7.10±0.90 µM on day seven (Table. S3), supporting the assumption that MGO has good tryptophan-trapping capacity. Indeed, the affinity of MGO to tryptophan also noticeably increased as tryptophan concentrations (0.1, 0.4, and 1.0 mM) increased (Fig. 5D). Given that fewer amino acids are known to be associated with depression and anxiety, the MGO-induced depression-like behavior in mice might be due to tryptophan depletion by MGO.

**Fig. 5:**
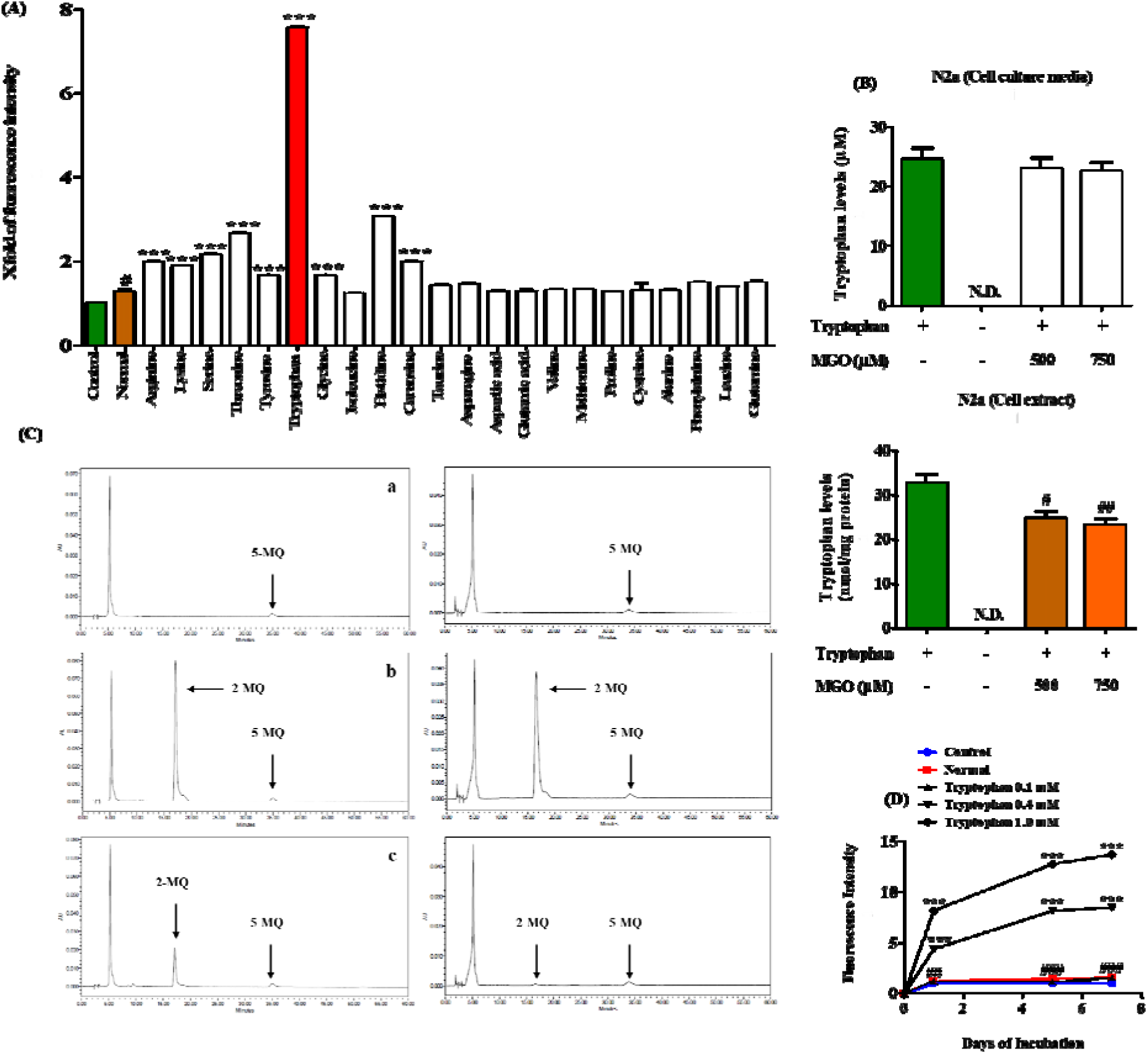
Affinity of MGO towards amino acids and trapping-ability towards tryptophan. (A) An affinity assay was performed to analyze the reaction between MGO and amino acids. The amino acids (1 mM) were incubated with MGO (10 mM) for 7 d. The affinity value of the MGO amino acid complex was evaluated using fluorescence at excitation/emission wavelengths of 355/460 nm. All data have been represented as mean ± SEM. n=3 (^#^*p*<0.05 *vs.* control; PBS and \*\*\**p*<0.001 *vs.* normal; PBS+MGO 10 mM). (B) N2a cells were seeded in a 60 φ dish, and extracted with ice-cold 50% methanol/50% PIPES-EDTA at -20°C for 5 to 10 min. Then, the samples were injected into the HPLC system. Tryptophan levels were detected using HPLC. All data have been represented as mean ± SEM. n=3 (^#^*p*<0.05, ^##^*p*<0.01 *vs* Tryptophan (+) medium treatment). (C) The trapping-ability of MGO towards tryptophan was analyzed using an MGO scavenger assay. HPLC chromatograms of tryptophan co-incubation with MGO (1:1) for 1 d and 7 d. (a) PBS (1 d); (b) PBS + 1 mM MGO (1 d); (c) PBS + 1 mM MGO + 1 mM tryptophan (1 d); (d) PBS (7 d); (e) PBS + 1 mM MGO (7 d); (f) PBS + 1 mM MGO + 1 mM tryptophan (7 d). n=3. (D) The fluoresce intensity upon the interaction of different concentrations (0.1, 0.4, and 1.0 mM) of tryptophan and 10 mM MGO were measured using an MGO-affinity assay. To confirm the MGO-reacting ability to tryptophan, tryptophan was incubated with MGO for 7 d. The affinity of MGO-tryptophan was measured in terms of fluorescence at excitation/emission wavelengths of 355/460 nm. All data have been represented as mean ± SEM. n=3 (^##^*p*<0.01, ^###^*p*<0.001 *vs.* PBS, and \*\*\**p*<0.001 *vs.* PBS+ MGO 10 mM).

### Tryptophan depletion by MGO induced cell apoptosis and damaged neuronal growth in N2a cells

To identify whether MGO-caused tryptophan deficits are responsible for the induction of apoptosis and damage to neuron structure, we performed Fluorescence-Activated Cell Sorting (FACS)/western blot analysis and Golgi staining assay, respectively. We treated cells with or without a tryptophan-containing medium in the presence or absence of MGO. Treatment with only tryptophan-free media resulted in cell toxicity and apoptosis in N2a cells. Similarly, treatment with MGO and tryptophan-containing media also induced cell toxicity and apoptosis in N2a cells (Fig. 6A-C). Western blot experiments with the same optimized conditions showed more induction of apoptosis in these conditions than that in the control. This was further supported by the higher Bax protein expression levels upon treatment with MGO (250 and 500 µM) + tryptophan-containing media than that upon treatment with or without only tryptophan-containing media in N2a cells (Fig. 6D-E). On the other hand, reduction in Bcl-2 protein expression level was more visible upon treatment with MGO (250 and 500 µM) + tryptophan-containing media than that with or without only tryptophan-containing media, thus resulting in an increased Bax/Bcl-2 ratio in N2a cells. A similar effect was observed in the case of Glyoxalase-I (GLO-I) and Glyoxalase-II (GLO-II) expression levels in N2a cells. Additionally, we examined lactate dehydrogenase (LDH) production under the same experimental conditions, where treatment with a tryptophan-free medium increased LDH production significantly as compared to that seen in the control, which was further reduced upon treatment with tryptophan-containing medium + MGO (250 and 500 µM). As D-lactate is the detoxified product of MGO, we analyzed D-lactate concentration as well, and the results showed that treatment with tryptophan-containing medium + MGO (250 and 500 µM) elevated D-lactate levels as compared to both the controls (Fig. S2A, B). Next, we used primary hippocampal cells to evaluate the effects of tryptophan deficiency and MGO using WST-1 assay and measurement of spine density. The WST-1 experiment revealed cell toxicity in the tryptophan-free medium + MGO-treated groups, as compared to the control group (Fig. 6F). Likewise, staining analysis revealed that the neuron spine density was different in each group. Tryptophan-deficient medium + MGO (500 and 750 µM) robustly decreased dendritic spine numbers as compared to those in the control (Fig. 6G and H). We also examined the effects of MGO + tryptophan treatment on neurite outgrowth in N2a cells. Treatment with tryptophan-deficient medium and MGO + tryptophan-containing medium markedly decreased neurite length, branch points, and cell body cluster, as compared to the control in N2a cells (Fig. S2C, D). From these results, it is clear that MGO facilitated apoptosis, and the reduction in neurite outgrowth and dendritic spine number was due to tryptophan deficiency.

**Fig. 6:**
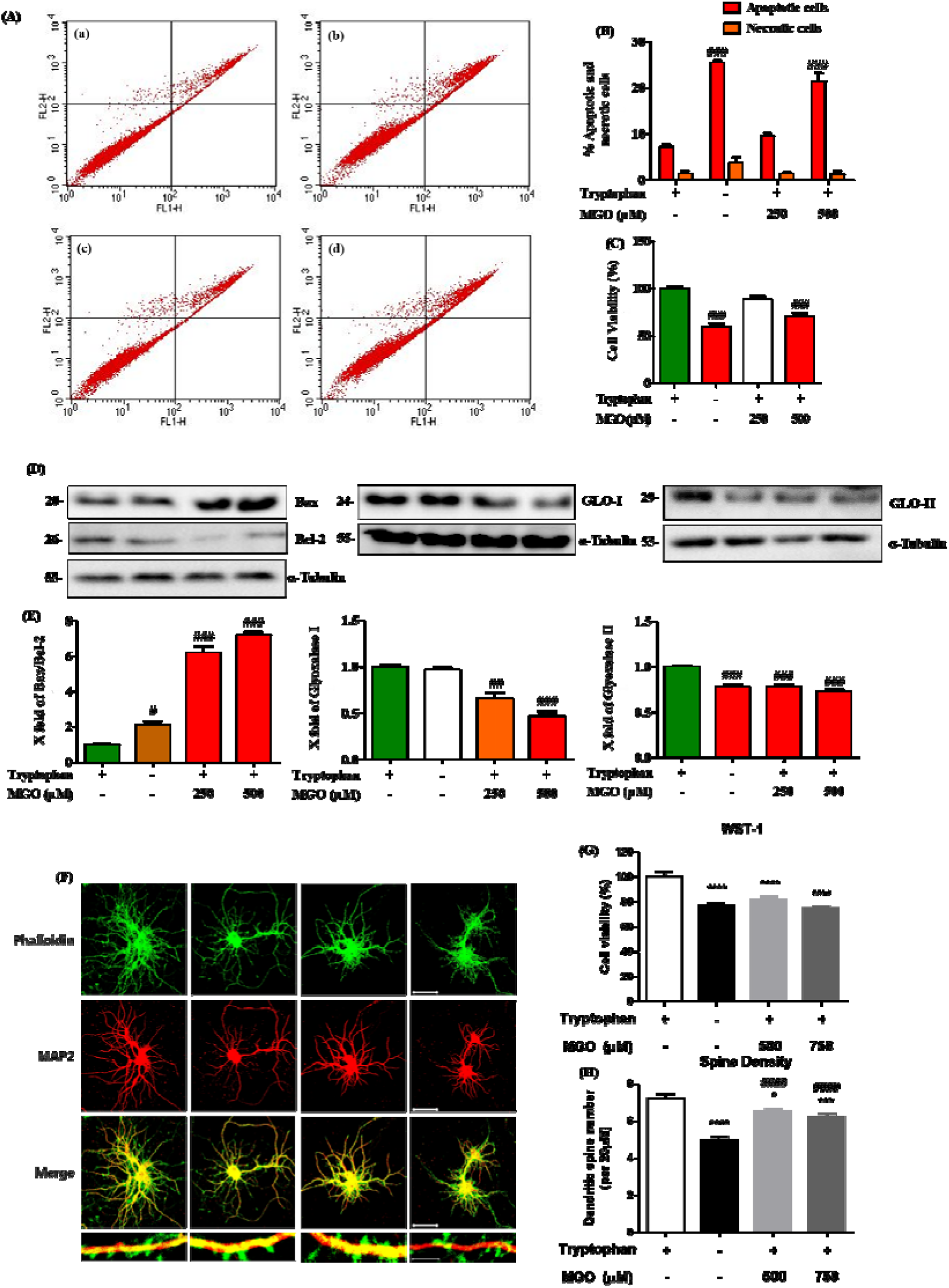
MGO and tryptophan deficiency-induced apoptosis and impaired neuronal development in primary neuron cells. (A) Cells were treated with different concentrations (250 and 500 μM) of MGO and tryptophan-free medium for 24 h. The necrotic and apoptotic cells were detected using flow cytometry with Annexin V-FITC and PI staining. (a) Tryptophan (+) medium; (b) Tryptophan free medium; (c) MGO 250 μM; (d) MGO 500 μM. (B) Percentage of cells in different phases: viable, early-stage apoptotic, late-stage apoptotic, and necrotic cells. Data have been represented as mean ± SEM. n=3 (^###^*p*<0.001 *vs.* Tryptophan (+) medium treatment). (C) MGO and tryptophan-free medium-induced cytotoxicity were measured using MTT assay in N2a cells. Data have been represented as mean ± SEM. n=3 (^###^*p*<0.001 *vs.* Tryptophan (+) medium treatment). (D) Cells were treated with different concentrations (500 and 500 μM) of MGO and tryptophan-free medium for 24 h. The protein expression levels of Bax, Bcl-2, GLO-I, GLO-II, and α-tubulin were detected using western blots. (E) Densitometry data for Bax, Bcl-2, GLO-I, and GLO-II were quantified using the Image Lab analysis tool. All data have been represented as mean ± SEM. n=3 (^#^*p*<0.05 and ^###^*p*<0.001 *vs.* Tryptophan (+) medium treatment). (F) Immunofluorescent staining of primary hippocampal neurons. The cells were treated with different concentrations (500 and 750 μM) of MGO and tryptophan-free medium for 24 h. The cells were fixed with DIV14 and immunostained. Representative images from three independent experiments. Scale bar: 100 µM. (G) WST-1 reagent was used to examine the cell viability of primary hippocampal neurons. Dendritic spine density was counted from neuronal dendritic segments. All data have been represented as mean ± SEM. n=3 (^###^*p*<0.001 and \*\*\**p*<0.001 *vs.* Tryptophan (+) medium treatment, \**p*<0.001 *vs.* Tryptophan free medium treatment).

### MGO-mediated tryptophan depletion altered the expression of depression-related genes in neuron cells

We performed microarray analysis to address any changes in the RNA expression levels of genes related to depression in MGO + tryptophan-containing medium- and tryptophan-free medium-treated cells. A hierarchical clustering system was applied to permit visual patterns of gene expression, as shown in Fig. 7A. We used four different groups of samples, including those treated with media with tryptophan (Normal), media without tryptophan (Control), and MGO + tryptophan-containing medium at concentrations of 250 µM and 500 µM. For visual inspection, a cluster tree was used, which indicated the gene expression patterns of the different treatment groups. Gene fold-change values in the presence of a particular treatment group are summarized in supplementary material (Table. S4). The tree cluster revealed a distinct pattern of gene expression in the treated and untreated groups (Fig. 7A). Surprisingly, MGO (500 µM) treatment dramatically downregulated the expression levels of protein tyrosine phosphatase receptor type T (Ptprt), protein-arginine deiminase type-2 (Padi2), TPH1, and TPH2 genes as compared to that in the normal control in N2a cells (Fig. 7B), while tryptophan-free medium significantly also decreased Padi2 but increased the mRNA levels of TPH2 and Ptprt. We also investigated the protein expression levels of these genes using western blot imaging. Data showed that the protein expression levels of Ptprt, Padi2, TPH1, and TPH2 decreased similarly to their mRNA levels in N2a cells. Notably, the downregulating effects of 500 µM MGO on mRNA and protein expression levels of Ptprt, Padi2, TPH1, and TPH2 were more prominent compared to tryptophan-free medium-treated cells (Fig. 7C-D). Additionally, we also checked tyrosine hydroxylase (TH) and synapsin-I protein expression levels under the same conditions, as they have flagship roles in neurotransmitter synthesis and synapse formation, respectively. Consistent with our previous results, 500 µM MGO significantly reduced TH and synapsin-I protein expression, although ptprt and TH expression started to increase upon treatment with high-concentration MGO (750 µM). Since ptprt is one of the best-characterized proteins due to its role in synaptic formation and neuronal development that are related to depression [53], we confirmed its expression pattern in the different treatment groups using immunofluorescence analysis. We found that treatment with MGO (250 and 500 µM) decreased fluorescence intensities in the presence or absence of tryptophan in N2a cells (Fig. S3A, B). Together, these data revealed that MGO triggered downscaling of synapse or neuron development-related genes/proteins, possibly by generating tryptophan deficiency.

**Fig. 7:**
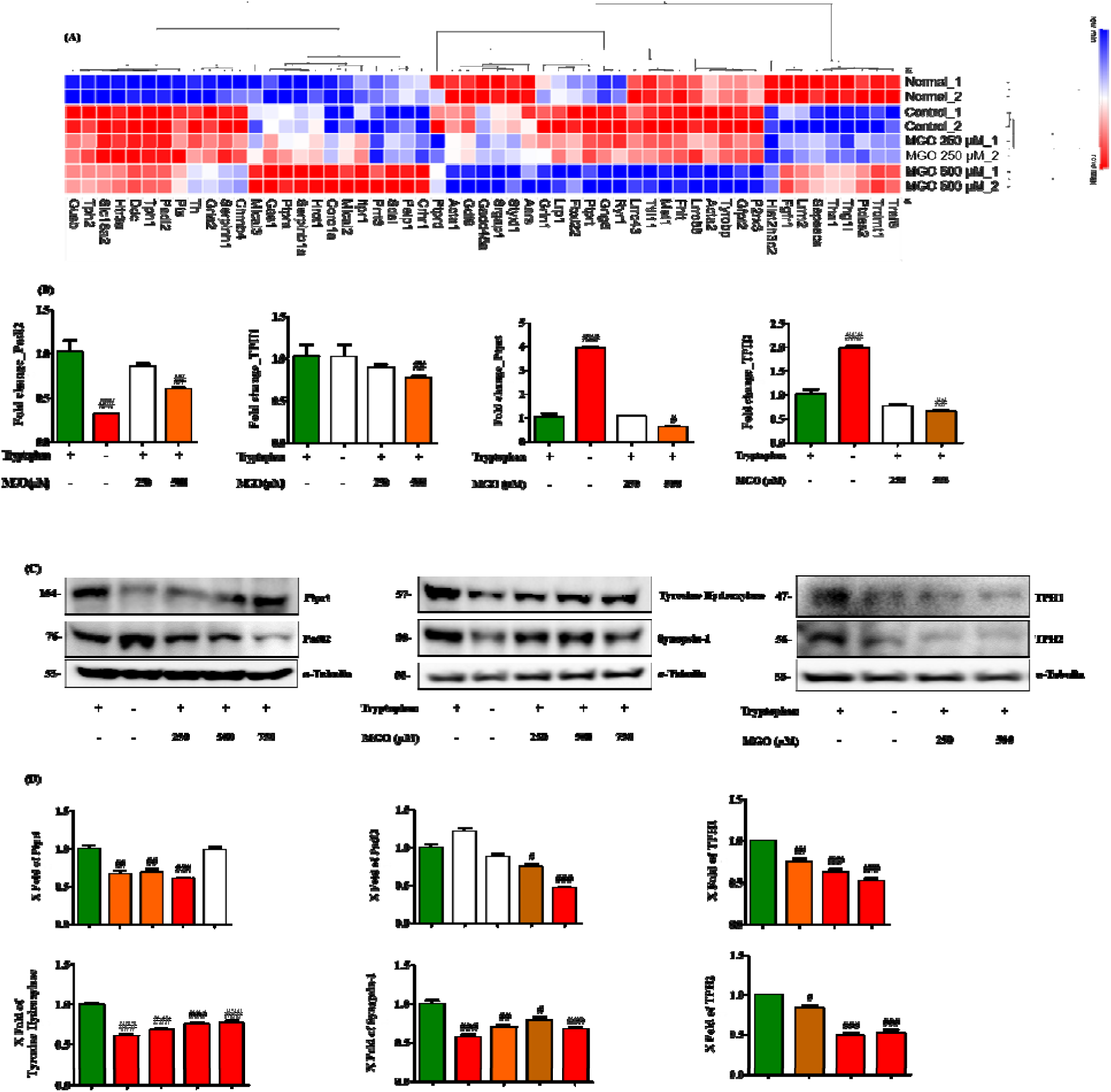
MGO and tryptophan deficiency up or downregulated the functional neuronal makers in N2a cells. (A) mRNA up- and downregulation, as presented using a Hierarchic clustering heat map. (B) Cells were treated with different concentrations (250 and 500 μM) of MGO and tryptophan-free medium for 24 h. MGO and tryptophan-free medium-induced mRNA expression were measured using quantitative real-time qPCR (RT-qPCR) in N2a cells. (C) N2a cells were treated with different concentrations (250, 500, and 750 μM) of MGO and tryptophan-free medium for 24 h. The protein expression levels of Ptprt, Padi2, tyrosine hydroxylase, synapsin-1, β-III-tubulin, and α-tubulin were evaluated using western blots with the appropriate antibodies. (D) The densitometry data of Ptprt, Padi2, tyrosine hydroxylase, synapsin-1, β-III-tubulin, and α-tubulin were determined using the Image Lab analysis tool. All data have been represented as mean ± SEM. n=3 (^#^*p*<0.05, ^##^*p*<0.01, and ^###^*p*<0.001 *vs.* Tryptophan (+) medium treatment).

### Tryptophan treatment rescued MGO induced depression and IBS in ICR mice

To finally demonstrate that MGO induced depression is mediated through tryptophan depletion, we exogenously administrated tryptophan at a concentration of 40 mg/kg by rectal injection in ICR mice in the presence or absence of MGO. Surprisingly, tryptophan treatment significantly reversed mice staying time in the central zone in the open field test, which was reduced by MGO (Fig. 8A, D). In the tail suspension and forced swimming test, the immobility time was prolonged with the MGO treatment mice group as compared to the control group, but the immobility time was reduced in tryptophan treatment mice (Fig. 8B and C). To support all these tests results, we also checked the effects of tryptophan on neurogenesis, tryptophan metabolism, and glyoxalase system-related proteins like synapsin-1, PTPRT, TPH-2, GLO-I, and GLO-II in mice brain tissue. In consistence with behavioral test results, tryptophan treatment dramatically increased synapsin-1, PTPRT and TPH-2 expression levels in the brain as compared to MGO treated depressive mice. Moreover, GLO-II and GLO-I protein expression levels significantly increased by tryptophan in MGO exposed depressive mice, suggesting its MGO scavenging efficacy (Fig. 8E). Further, we decided to evaluate tryptophan treatment effects in MGO induced IBS model, and the results were noticeable. Tryptophan exclusively enlarged the colon length of MGO induced depressive mice as compared to only MGO treated depressive mice’s colon length (Fig. 8G). Moreover, tryptophan treatment reversed the MGO effects on ZO-1 and Ki67 in mice colon tissue (Fig. S4C, D). Additionally, it also reduced pro-inflammatory cytokines IL-6 and IL-1β secretion in the plasma as compared to MGO treated group (Fig. 8H and I). These results suggest that the presence of tryptophan might protect mice from depression and IBS through its MGO trapping capacity and allow homeostasis conditions for tryptophan metabolism in the brain and reduce inflammation in the large intestine, respectively.

**Fig. 8:**
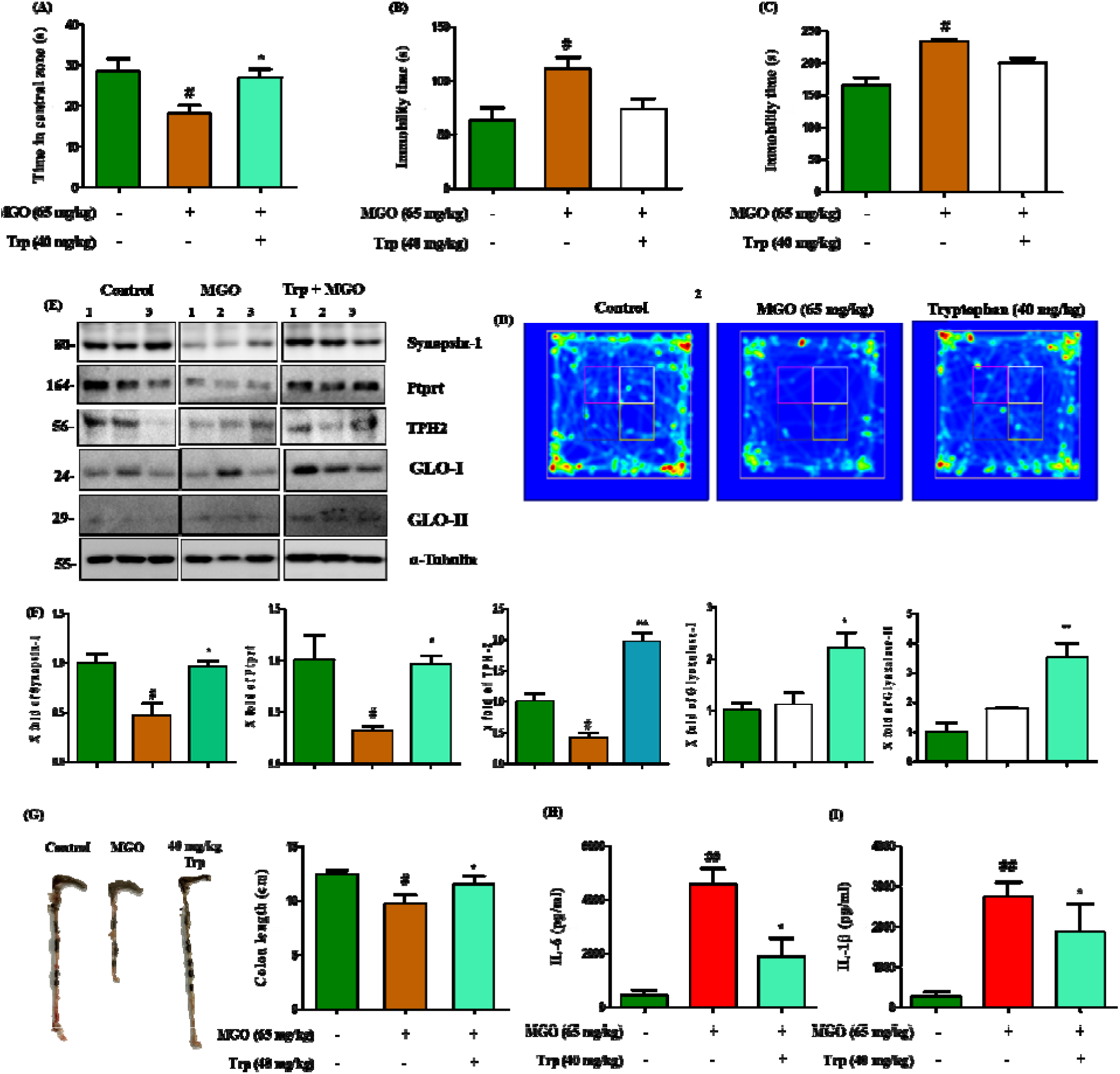
Tryptophan rescued MGO induced depression and IBS in mice. (A) OFT was done in mice after 3 weeks of MGO and tryptophan (Trp) administration by the rectal route. Mice were exposed to the open field for 8 min, to examine the time spent in the central zone. Data are presented as mean ± SEM. n=6 (^#^*p*<0.05 vs control group, \**p*<0.01 vs MGO treated group) (B, C) The immobility time of mice exposed to MGO and tryptophan in TST and FST. Data are presented as mean ± SEM. n=6 (^#^*p*<0.05 vs control group, \**p*<0.108 vs MGO treated group) (D) Visual recording of the OFT was carried out using SMART3.0 SUPER PACK. (E-F) Expression of synapsin-1, PTPRT, TPH-2, Glo-I, and Glo-II in mice brain tissue injected with MGO and Tryptophan. Data are presented as mean ± SEM. n=3 ((^#^*p*<0.05 vs control group, \**p*<0.05 vs MGO treated group, \*\**p*<0.05 vs MGO treated group \*\*\**p*<0.05 vs MGO treated group). (G) The normal control group received only 30% v/v glycerol in PBS, while the two other groups received tryptophan (Diluted in PBS) in the presence or absence of MGO. After sacrifice, the colon length was measured using a ruler (in cm). Data have been represented as mean ± SEM. n=6 (^#^*p*<0.05 *vs.* control, \**p*<0.01 *vs.* MGO treated group). (H, I) The secretion of IL-6 and IL-1β cytokines in the plasma was evaluated using the respective ELISA assay kit. Data have been represented as mean ± SEM. n=6 (^##^*p*<0.001 *vs* control and \**p*<0.01 *vs.* MGO treated group).

## Discussion

Numerous studies have found a relationship between hyperglycemia and brain dysfunction, particularly depression, and behavioral flexibility in diabetic patients [54, 55, 56]. In this context, data suggest that high levels of glucose are greatly responsible for brain damage, and there are many arguments about whether a high-carbohydrate diet is the cause of depression or indulged in depression. If hyperglycemia persists due to various causes, eventually, this elevated glucose condition produces different dicarbonyls such as MGO, GO, glyceraldehyde (GA), and glycolaldehyde (GC) as a byproduct of the glycolysis process; MGO is the most reactive among them [57]. Moreover, consistent studies have revealed that MGO and its advanced glycation end-products hinder the cognitive function of mice and humans [58, 59]. In addition, a large body of reports has demonstrated the involvement of MGO-induced protein glycation in impaired cognitive function [60, 61]. However, the gut microbiome produces different kinds of metabolites, such as short-chain fatty acids, bile acids, and MGO, of which, MGO is also a highly reactive component [62]. Moreover, one study found that MGO may have some correlation with IBS, although the actual mechanism is not yet clear. Thus, MGO’s function in IBS and its link to brain depression have not been properly studied to date. Nonetheless, tryptophan production and metabolism in the gut and the role of gut microbiota in tryptophan-mediated brain dysfunction have been recognized for a long time. 5-HT is a tryptophan metabolic product that is generated in the brain; however, more than 90% of its production occurs in the intestine [63]. Less availability of neurotransmitters such as 5-HT has been implicated to have a role in the gut-brain axis in depression and anxiety [64]. Moreover, emerging evidence suggests that many diseases, such as IBS, colitis, non-alcoholic fatty liver disease, and particularly anxiety and depression, are markedly affected by tryptophan metabolism [65]. However, since MGO is abundant in the body in hyperglycemic conditions and produced by the intestinal gut microbiota, and tryptophan is also available in the intestine, we hypothesized that MGO might be linked to tryptophan, in the context of it leading to depression in the brain and damage to brain cells and IBS in the intestine.

In this study, to identify the direct role of high dose MGO in the gut, we used the rectal route for the administration of MGO to mice, as this model has been previously reported as a diabetic model [66]. We repeatedly performed depression-like behavioral tests such as OFT, a TST, and an FST to verify our hypothesis in mice. We found that as the MGO dose increased, there was a reduction in the time that the mice spent in the central area and an increase in their immobility (Fig. 1A-D). It is known that 5-HT, EP, NE, and DA are the representative neurotransmitters involved in depression, anxiety, and other cognitive impairments, although the last two have been less studied in this context [67, 68, 69]. In our experimental model, MGO displayed remarkable effects on decreasing the levels of 5-HT, EP, NE, and DA in mouse brains (Fig. 1E-H). A study conducted on a distinct part of cognitive function in patients with depression revealed a greater loss of hippocampal activity [70]. The hippocampus is an anatomically connected part of the brain that is involved with memory, learning, and depression behavior.

This system is composed of three components: the DG, subiculum, and CA fields, which are subdivided into four regions (CA1-CA4). The cortex region transfers information to the hippocampus via two pathways: one through the DG and CA3 areas, and the other through the subiculum and CA1. It is strongly believed that the hippocampus allows long-term memory in a process termed LTP, after receiving and consolidating the information from the entorhinal cortex. LTP is induced when neurotransmitter release occurs 5-15 milliseconds before a back pro-purgatory action potential. Therefore, it seems that high dose MGO induced LTP impairment; however, one research group reported that when they engaged more than 2000 subjects to study the hippocampal areas of patients with depressive disorder, they found that fewer hippocampal areas in patients with depression [4, 71]. Therefore, we investigated whether the effects of MGO in inducing depression-like behavior in mice (as observed in the first part of our study) are mediated through damaging the hippocampus. Consistent with previous findings, our results indicated that MGO sharply reduced the number of cells in the DG, CA3, and CA1 areas of the mouse hippocampus (Fig. 2A-D). Moreover, the LTP of the hippocampus also dramatically declined upon MGO treatment (Fig. 2E-F), supporting pathological changes in the hippocampi of mice. Therefore, our data propose that MGO may be a new molecular mediator of glucocorticoid receptor activity. But more in-depth research is needed to elucidate this suggestion.

Tryptophan is the main substrate for 5-HT synthesis, and 5-HT continuously plays a role in the brain, in terms of depressive and behavioral responses [72]. Biosynthesis of 5-HT is a rate-limiting step, which is generally catalyzed by the enzyme TPH. Given this, we analyzed the effects of MGO on tryptophan, and its derivatives; for instance, 5-HTP and 5-HT have a critical role in depression and anxiety, and data from our study showed decreased levels of tryptophan, 5-HTP, and 5-HT in the mouse plasma (Fig. 3A-C). TPH can be found in two of its isoforms, TPH1, and TPH2, and it is suggested that the TPH2 form of hydroxylase is rigorously expressed in the brain and periphery. Consistent with this, we can suggest that less availability of this enzyme in the human brain may be strongly related to depression-like behavior [29, 30]. In this context, we observed the effects of MGO treatment on TPH1 and TPH2 through histochemical analysis of brain sections. Results from this assessment showed that MGO significantly reduced TPH1 and TPH2 levels in the CA1, CA3, and cortex regions (Fig. 3D-J), suggesting further reduced 5-HT levels in that area of the brain. After finding the damaging effect of MGO on tryptophan and its metabolically-related enzymes and neurotransmitters (which are truly associated with depression) in brain sections, we investigated the effect of MGO on these factors in the intestine. This was based on a previous demonstration that the gut-brain axis is an established model because of its functional interconnection [73]. MGO is abundantly found in the intestine, and TPH1 also is highly expressed in the intestine. Surprisingly, we found that MGO decreased TPH1 levels in colon tissue and colon length and increased inflammation (Fig. 4A-J), while the body weight remained unchanged throughout the treatment period, which was a clear indication of IBS.

Based on the *in vivo* results, we hypothesized that MGO might have direct crosstalk with tryptophan in the case of depression and behavioral changes in mice and dysfunctional syndrome in the intestine like IBS. To clarify our assumption, we initially performed an MGO affinity assay to examine whether MGO has a high affinity for tryptophan or other amino acids (Fig 5A). Since their metabolism and availability are attractive potential factors in the maintenance of neurotransmitter generation, surprisingly, we noticed that MGO showed the highest affinity for tryptophan, but not to other amino acids. To clarify these findings, we also detected the residual tryptophan level by the tryptophan-trapping ability of MGO through HPLC analysis (Fig 5C).

For identifying associations, we checked MGO toxicity in different types of brain cells; among them, neuron cells (N2a) showed the highest sensitivity to MGO. In agreement with this result, we gave cells treatment without tryptophan and with the combination of MGO and tryptophan (MGO+Tryp). Upon treatment without tryptophan, we observed apoptosis induction and reduced neurite growth in N2a cells. However, greater effects on apoptosis-related proteins and neurite outgrowth and its related proteins were discovered when cells were treated with the combination of MGO and tryptophan (Fig 6A-D). Of specific relevance, combined treatment with MGO+Tryp diminished the levels of the MGO-detoxifying enzymes, GLO I and GLO II, more prominently, as compared to treatment without tryptophan in neuronal cells.

Our previous results, however, suggest that the MGO-trapping capacity of tryptophan or deactivating tryptophan activity might have fatal effects on neuronal growth and development-regulatory markers. To investigate this, we performed microarray analysis to identify any changes upon MGO +Tryp treatment in N2a cells and found significant changes in some genes that are responsible for synaptic function and neuronal development. MGO treatment remarkably altered levels of Ptprt, Padi2, TH, and synapsin-1 in the presence of tryptophan (Fig 7), as they play a critical role in tryptophan metabolism and neuronal development in the brain and intestine [53, 74, 75]. It has been demonstrated that Ptprt knockdown induces depression-like behavior in mice through defective synaptic function and abnormal neurogenesis [53]. Moreover, the importance of Padi2 and synapsin-1 in oligodendrocyte differentiation and myelination has been reported [74, 76]. The above results indicate that the MGO-induced depression-like effects in mice and defective neuron cell growth might be through the regulation of these genes in mice.

To clarify our hypothesis that MGO mediated depression and IBS in mice by the diminished tryptophan levels, we treated depressive mice with tryptophan, and the outcomes were exclusionary. In the OFT test, Tryptophan administered mice showed a marked tendency in the number of line crossing and increased spending time in the center. Similarly, the immobility time was reduced by the tryptophan treated mice group compared to the depressive model group. Interestingly, tryptophan facilitates the increasing expression of neurogenesis and tryptophan metabolism-related proteins even in MGO-induced depressive mice. It is noteworthy that Tryptophan accelerated MGO detoxification factors like GLO I and GLO II, indicating tryptophan engulfing characteristics to MGO may promote to show tryptophan anti-depression effects. In the IBS model, tryptophan dramatically increased the colon length of mice which was reduced by MGO and declined inflammation as well.

## Conclusions

In summary, MGO accelerated the reduction of tryptophan metabolism-related factors TPH1, and TPH2 levels in the mouse plasma, and the hippocampal area, while it also reduced the levels of neurotransmitters exclusively linked to depression-like behavior. Interestingly, it was shown similar phenomena in the mice intestine, including induced reduction of colon size and inflammatory bowel syndrome. Treatment with MGO + Tryptophan displayed almost similar effects as that exhibited upon tryptophan-null treatment in neuronal cells, such as induction of cell toxicity, decrease TPH1, and TPH2 levels, and inhibition of dendritic spine density and neuronal outgrowth. Importantly, exogenous tryptophan administration protects mice from MGO-induced depression-like behavior and inflammatory bowel syndrome (Fig. 9). However, our study strongly demonstrated that it is important to raise the level of tryptophan to prevent MGO induced neuronal dysfunction.

**Fig. 9:**
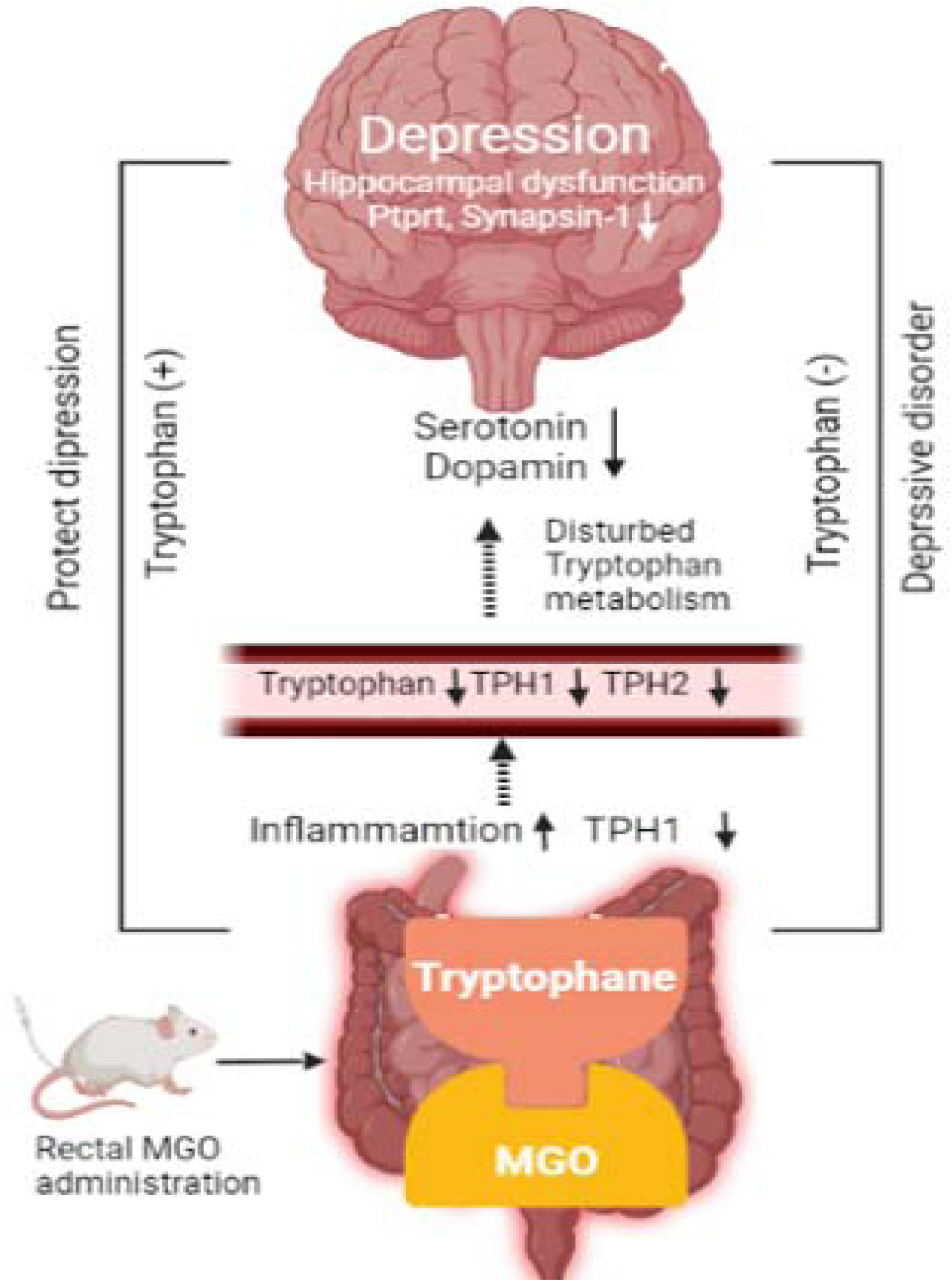
Schematic diagram of the mechanism of MGO induced depression and neuronal dysfunction in mice via tryptophan depletion. MGO decreased the tryptophan metabolism marker in the colon and brain and reduced neurotransmitters and hippocampal cells. It also facilitated apoptosis in neuronal cell and inflammation in mice. Tryptophan treatment significantly improved MGO triggered dysfunction colon and brain.

## Availability of data and materials

This study includes no data deposited in external repositories. All data are included in this article and additional file 1 and are available upon request from the corresponding author.

## Acknowledgments

We are also thankful to Myoung Gyu Park, MetaCen Therapeutics Co., Ltd for his valuable support.

## Funding

This research was supported by a grant from the Basic Science Research Program through the National Research Foundation of Korea (NRF), funded by the Ministry of Education (NRF-2018R1D1A1B07049500).

## Authors Contributions

Conceptualization, design, experiments: MS, LJH, KSY. Data curation: MS, LJH. Formal analysis: MS, LJH, SMH, HBK, HP, KAC, HE, and MSO. Drafting: M.S. Funding acquisition: KSY. Methodology: MS, LJH, HP, KAC. Supervision: KSY. Review, and editing: MS, and KSY.

## Competing interests

The authors declare no conflict of interest.

## Ethics approval and consent to participate

All animal experiments were performed in compliance with the ethical requirements of the laboratory research center, college of pharmacy, Gachon University, Korea, and the protocol was approved by the Gachon University Institutional Animal Care and Use Committee. This study does not contain any patient samples experiment.

## Consent for publication

Not applicable

## Abbreviations

DM: Diabetes mellitus
MGO: Methylglyoxal
GO: Glyoxal
TPH1: Tryptophan hydroxylase 1
TPH2: Tryptophan hydroxylase 2
CEL: Carboxymethyl-lysine
MGH1: Methylglyoxal-derived hydroimidazolones
5-HT: 5-hydroxytryptamine
5-HTTP: 5-hydroxytryptophan
DA: Dopamine
EP: Epinephrine
NE: Norepinephrine
DG: Dentate gyrus
CA3: Cornu ammonis 3
CA1: Cornu ammonis 3
Bax: BCL-2 Associated X protein
BcL-2: B-cell lymphoma 2
GLO-I: Glyoxalase-I
GLO-II: Glyoxalase-II
PTPRT: Protein tyrosine phosphatase receptor type T
Padi2: Protein-arginine deiminase type-2

## Figure legends

## Notes

### Competing Interest Statement

The authors have declared no competing interest.

