## Supplementary material for "Methylglyoxal is the main culprit to impairing neuronal function: mediated through tryptophan depletion": Supplimentary file.docx

^g^ MetaCen Therapeutics Company, Changnyong-daero 256 beon-gil, Yeongtong-gu, Suwon-si, Gyeonggi-do, 16229, Republic of Korea.

^h^ College of Pharmacy, Kyung Hee University, 26 Kyungheedae-ro Dongdaemun-gu. 02447, Seoul, Republic of Korea.

*Correspondence: Sun Yeou Kim

College of Pharmacy, Gachon University, #191, Hambakmoero, Yeonsu-gu, Incheon 21936, Republic of Korea.

**Supplementary methods**

**Material**

Glyoxal (GO), MGO, perchloric acid (PCA), *o*-phenylenediamine (*o*-PD), tryptophan, 5-hydroxytryptophan (5-HTP), 5-HT, DA, E, NE, 2,7-dichlorofluorescein diacetate (DCFH-DA), thiazolyl blue tetrazolium bromide (MTT), and anti-tyrosine hydroxylase antibody were obtained from Sigma-Aldrich (St. Louis, MO, USA). Annexin V-FITC Apoptosis Detection Kit (San Diego, CA, USA) was obtained from BD Biosciences Pharmingen (Franklin Lakes, NJ, USA). JC-10 mitochondrial assay kit and anti-protein tyrosine phosphatase receptor type T (PTPRT), anti-histone H3 (di methyl K9), anti-histone H3, anti-synapsin, and anti-tryptophan hydroxylase (TPH)2 antibodies were purchased from Abcam (Cambridge, UK). Antibodies against α-tubulin, histone H3K4me3, protein arginine deiminase type 2 (PADI2), and β-III-tubulin were purchased from Thermo Fisher Scientific (Waltham, MA, USA). Antibodies against sirtuin1, p38, p-p38, extracellular receptor kinase (ERK), p-ERK,c-jun N-terminal kinase JNK, and p-JNK were purchased from Cell Signaling Technology (Danvers, MA, USA). Antibodies against glyoxalase 1 (GLO-I), glyoxalase 2 (GLO-II), Nrf-2, Bcl-2, Bax, and p53 were purchased from Santa Cruz Biotechnology (Santa Cruz, CA, USA). 2-Methylquinoxaline (2-MQ) and 5-methylquinoxaline (5-MQ) were purchased from Tokyo Chemical Industry (Tokyo, Japan). Interleukin 1 beta (IL1β), IL-6, IL-10, and Tumor necrosis factor (TNF-α) ELISA kits were purchased from R&D Systems (Minneapolis, MN, USA). The NAD^+^/NADH assay kit was from Cell Biolabs Inc. (San Diego, CA, USA) and the EnzyChrom™ D-Lactate Assay Kit was from BioAssay Systems (EDLC-100, California USA).

**In vivo Methods**

**Open field test (OFT)**

The OFT was conducted in an open plastic box (45 × 45 × 45 mm), as reported by Hiroshi Ueno et al. with a bit change [38]. The zone of the box was divided into 24 grids of 11.25 × 11.25 cm. The mice were individually placed in the center of the open plastic box and allowed to freely observe, following which a video was recorded for 5 min. The open plastic box was cleaned with 70% ethanol. The time in the central zone(s) was analyzed using SMART3.0 SUPER PACK (Panlab; Harvard Apparatus, Barcelona, Spain).

**Tail suspension test (TST)**

TST was performed as mentioned by Kang et al with a bit of change [39]. The TST was conducted in a TST chamber (60 cm length, 60 cm height, 11.5 cm depth, and 15 cm width) and the mice were hunged by painless tape-based fixation. Before recording, all mice were pre-adapted to the TST chamber for 2 min. After the mice were individually placed in the TST chamber, the video was recorded for 4 min. Next, the immobility of the mice was analyzed using SMART3.0 SUPER PACK.

**Forced swimming test (FST)**

The FST was executed as reported by Kang et al with slight modifications [39]. It was performed in an FST chamber (50 cm height × 20 cm diameter) filled with water (at a level of 30 cm) set at room temperature (25±1°C). Before recording, all mice were pre-adapted to the FST chamber for 2 min and then placed in the FST chamber, following which the video was recorded for 4 min. The immobility of the mice was then evaluated using SMART3.0 SUPER PACK.

In vitro methods

**Enzyme-linked immunosorbent assay (ELISA)**

The treated or untreated conditioned plasma was used to measure the levels of secreted inflammatory mediators. Cytokine levels were quantified in the supernatants using respective ELISA kits (R&D Systems), according to the manufacturer’s instructions.

**D-Lactate assay**

D-Lactate production was evaluated using the EnzyChrom™ D-lactate assay kit (EDLC-100; BioAssay Systems), according to the manufacturer’s protocol. Briefly, CM from each well was transferred to a 96-well plate and mixed with an equal volume of reagent solution (60 µL assay buffer, 1 µL enzyme A, 1 µL enzyme B, 10 µL NAD, and 14 µL MTT). The D-lactate quantity was measured using a microplate reader at the wavelength of 565 nm.

**Primary hippocampal neuron culture**

Timed pregnant (TP) 17 days of SD rats were purchased from Koatech (Korea). Hippocampal tissue was dissociated out and 0.25% trypsin solution (Gibco) was used to enzymatically dissociate the tissue. The dissociated hippocampal tissue was washed using 1x HBSS (Gibco) and resuspended to neurobasal media containing 2% B27, 2-mM L-glutamine, and 1% Penicillin-Streptomycin (Gibco). Hippocampal neurons were seeded in 96 well plates or 18-mm coverslip pre-coated with 0.1 mg/ml Poly-L-lysine (sigma). Hippocampal neurons were plated on 96 well plates at a density of 2ｘ10^4^ cells/well or 18-mm coverslips at a density of 3ｘ10^4^ cells/coverslip, respectively. Culture media were changed and half-replaced every 3-4 days. Primary hippocampal neurons were maintained in a 5% CO^2^ humidified incubator at 37℃ for 14 days. At DIV 11-13, the media of primary hippocampal neurons was changed with media with or without tryptophan before treatment with 500 or 750 μM MGO.

**Cell viability assay for primary neuron cell**

WST-1 (Roche, Switzerland) measures the metabolic activity of viable cells. WST-1 reduction was determined according to the manufacturer’s instructions manual. After MGO was incubated for 24 hr, WST-1 reagent was added to the well. The primary hippocampal neurons were incubated at 37℃ in 5% CO_2_ for 2 hr. The absorbance of reaction media was measured at 450 nm using a vector X4 multi-label plate reader (Perkin Elmer).

**Western blot analysis**

N2a cells were seeded in a 60 φ dish, washed with cold PBS, and lysed in PRO-PREP™ protein extraction solution (iNtRON, Seoul, Korea) at -20°C for 24 h. After the cell lysates were separated by means of centrifugation, the protein concentration was determined using the Bradford assay. Proteins (30 µg) were separated using sodium dodecyl sulfate-polyacrylamide gel electrophoresis and transferred to Polyvinyleden fluoride membrane (PVDF) membranes using a Trans-Blot^®^ Turbo™ Blotting System. The membranes were then blocked using 5% skim milk for 1 h and incubated overnight at 4°C with primary antibodies. After overnight incubation, the membranes were washed and incubated with secondary antibodies at room temperature (25°C) for 1 h. The bands were detected using a ChemiDoc™ XRS+ imaging system (Bio-Rad, CA, USA).

**Supplementary figure legends**

Supplementary fig. 1: Effects of MGO on cell viability and migration in HIEC-6 cells.

(A) Treatment of HIEC-6 cells with MGO at different concentrations (250, 500, 750, and 1000 μM) for 24 h. Photomicrographs were investigated using phase-contrast microscopy (10× magnification). (B) Cell viability and migration rate-related factors (relative wound density, wound confluence, and wound width) decreased upon exposure to MGO for 24 h (500, 750, and 1000 μM) in HIEC-6 cells. All data have been represented as mean ± SEM. n=3 (^#^*p*<0.05, ^##^*p*<0.01, and ^###^*p*<0.001 *vs.* control).

**Supplementary fig.** **2: Effects of MGO and tryptophan deficiency on LDH production and D-lactate levels in N2a cells.** (A-B) Cells were treated with different concentrations (500 and 750 μM) of MGO and tryptophan-free medium for 24 h. Quantitative levels of LDH production and D-lactate were measured using respective assay kits in N2a cells. All data have been represented as mean ± SEM. n=3 (^###^*p*<0.001 *vs.* Tryptophan (+) medium treatment). (C) N2a cells were seeded into collagen-coated 6-well plates in the presence or absence of different concentrations (500 and 500 μM) of MGO and tryptophan-free medium for 24 h. Quantitative analyses of neurite outgrowth were conducted by calculating the number of neurite lengths from randomly selected fields per well. All images were captured at 20× magnification. Scale bar: 200 µm. n=3. (D) Neurite length, neurite branch points, and cell-body clusters were measured using Phase NeuroTrack software and plotted over time. All data have been represented as mean ± SEM. n=3 (^#^*p*<0.05 and ^###^*p*<0.001 *vs.* Tryptophan (+) medium treatment).

**Supplementary** **fig. 3: Assessment of effects of MGO and tryptophan deficiency on Ptprt fluorescence intensities in N2a cells using immunofluorescence analysis.**

(A) N2a cells were treated with different concentrations (500 and 750 μM) of MGO and tryptophan-free medium for 24 h. The cells stained with Hoechst 33342 (blue) and Ptprt (red) were observed using confocal microscopy. (B) The quantitative measurements of Ptprt fluorescence intensity were determined using NIS-Elements imaging software. Scale bar: 25 μm. All data have been represented as mean ± SEM. n=3 (^#^*p*<0.05 and ^###^*p*<0.001 *vs.* Tryptophan (+) medium treatment).

**Supplementary** **fig. 4: Effects of MGO and tryptophan on proliferation and tight junction markers in mice colon tissue.**

(A, C) Immunohistochemical (IHC) analysis of colon sections was performed. Representative images were stained for ZO-1 and Ki67 in the colon tissues. Scale bar: 200 µm. (B, D) Quantitative measurements of ZO-1 and Ki67 were conducted by calculating the expression in the colon from selected fields per image. The values were calculated using ImageJ software. Data have been represented as mean ± SEM. n=3 (^##^*p*<0.01 *vs.* control, and ^*^*p*<0.04 and ^**^*p*<0.01 *vs.* MGO treatment).

**Figure S1**

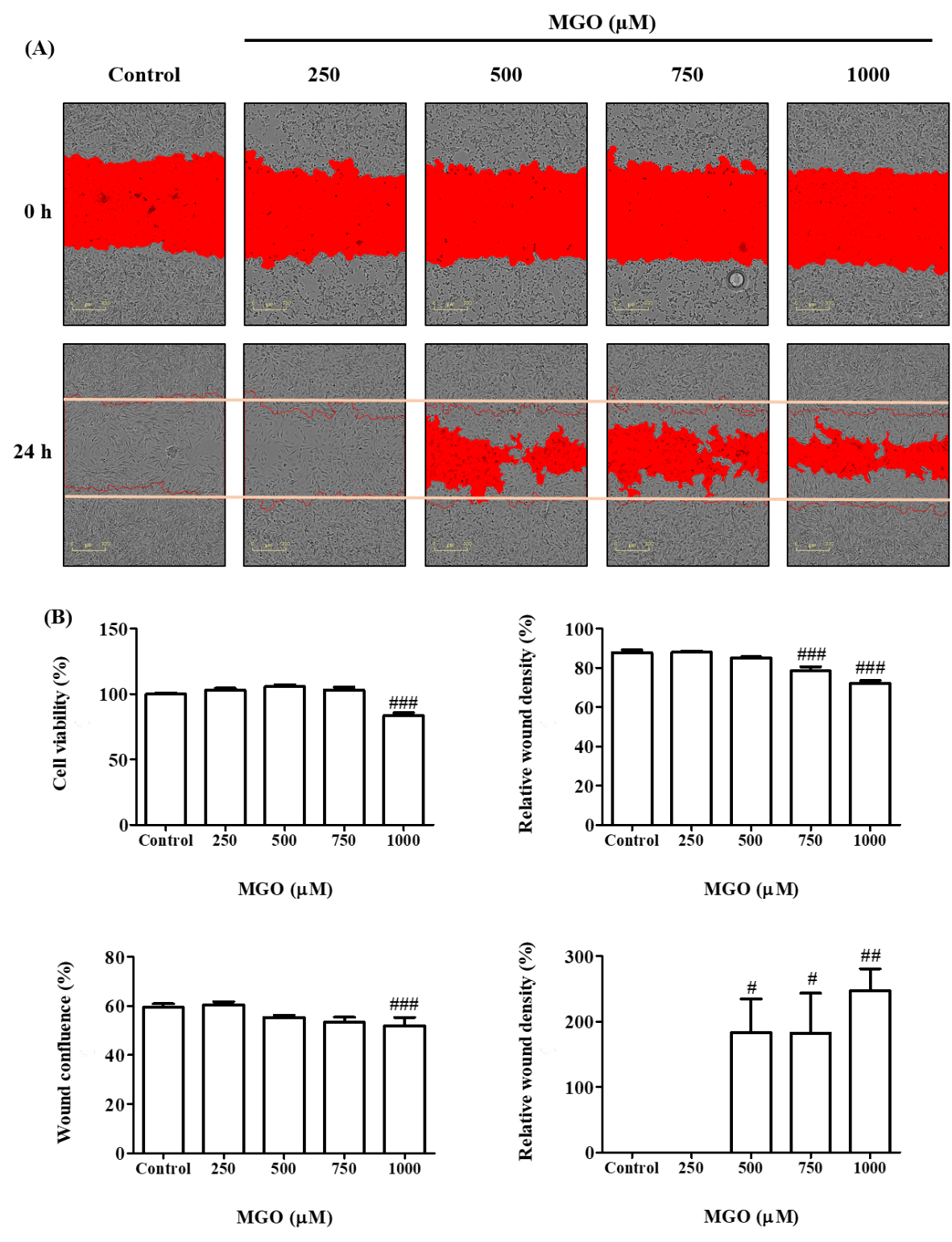

**Figure S2**

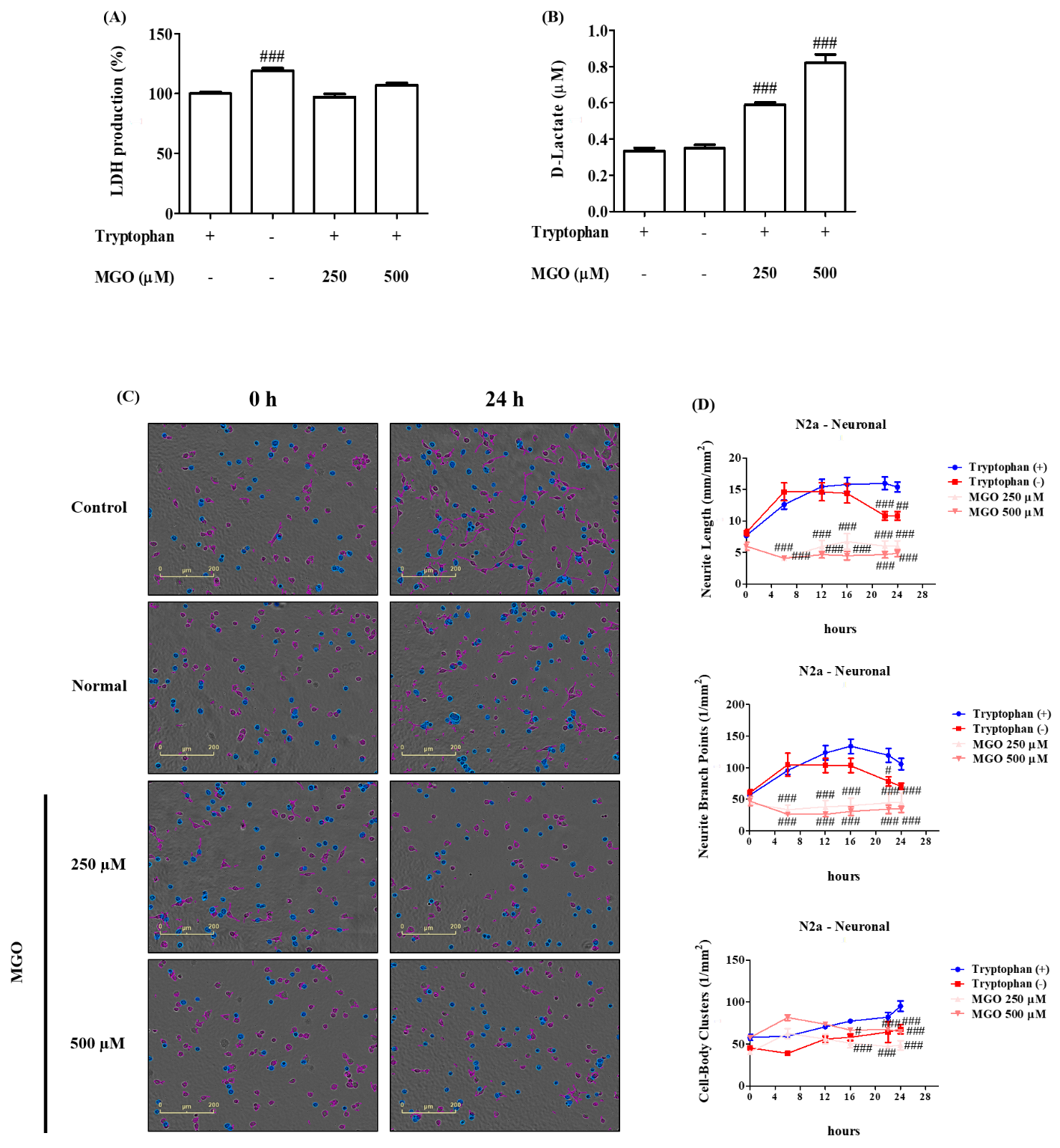

**Figure S3**

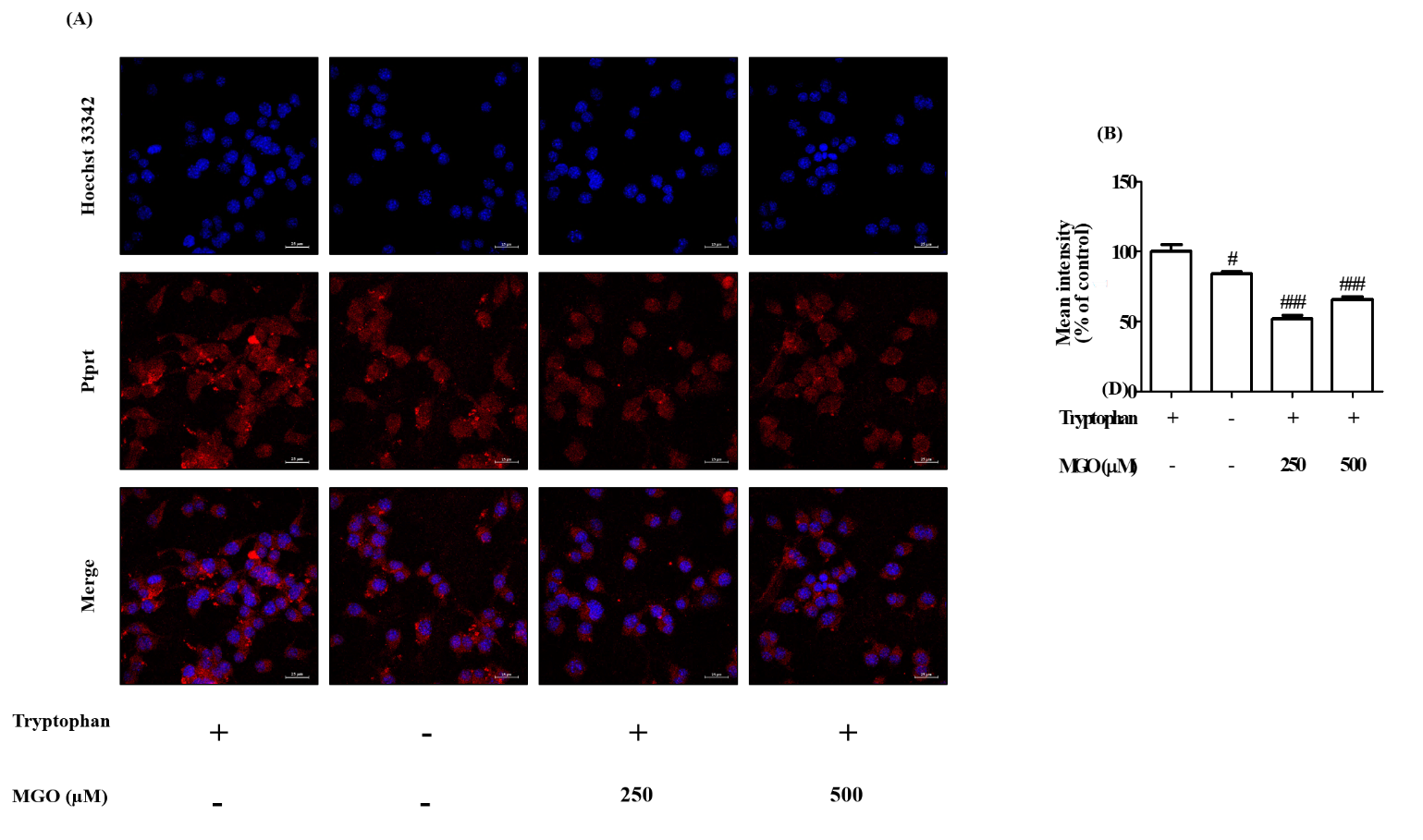

**Figure S4**

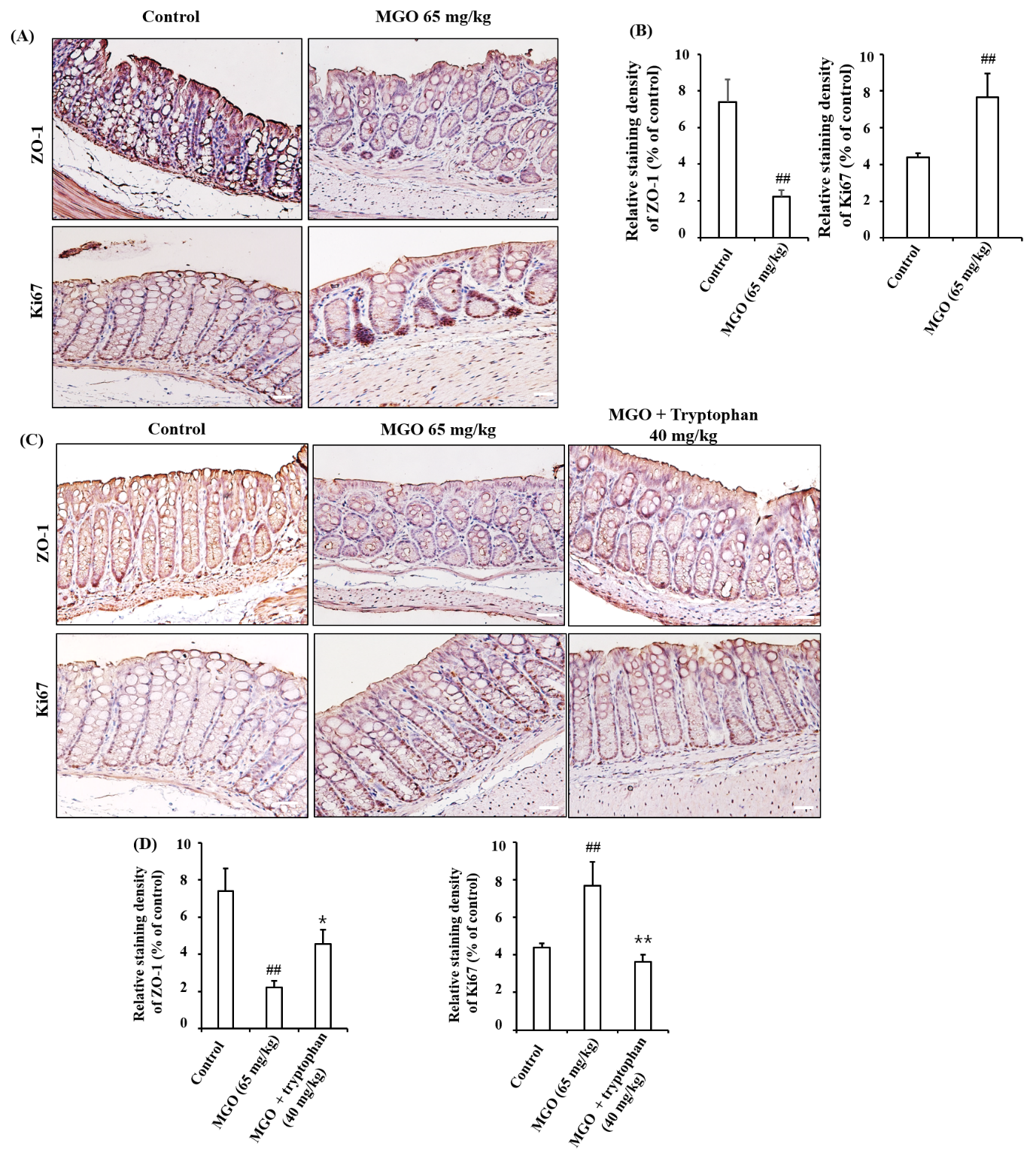

Table S1. Lists of primers used in qRT-PCR analysis.

| Gene | Direction | Sequence (5’ to 3’) |
| --- | --- | --- |
| TPH1 | Forward | TTCACCATGATTGAAGACAAC |
|  | Reverse | TCCGACTTCATTCTCCAAGG |
| TPH2 | Forward | CCATCGGAGAATTGAAGCAT |
|  | Reverse | TTCAATGCTCTGCGTGTAGG |
| Ptprt | Forward | ACCTGCTTCAACACATCACCCAGA |
|  | Reverse | TTCATCTTCCTTGGCTGTGTCCCA |
| Padi2 | Forward | GTAGGCCACGTCGATGAGTT |
|  | Reverse | TCCCAGGCCCTTGAACATAAC |
| GAPDH | Forward | TGCACCACCAACTGCTTAG |
|  | Reverse | GGATGCAGAGAAGATGTTC |

Table S2. Levels of tryptophan in the cell extract.

| Sample | Cell culture medium  (µM) | Cell extract  (nmol/mg protein) |
| --- | --- | --- |
| Control | 24.58 ± 1.92 | 32.84 ± 1.88 |
| Normal | N.D. | N.D. |
| MGO 250 µM | 23.11 ± 1.79 | 24.75 ± 1.57^#^ |
| MGO 500 µM | 22.63 ± 1.41 | 23.33 ± 1.27^##^ |

All data are shown as means ± SEM. N = 3 (#p < 0.05, ##p < 0.01 vs. Control).

Table S3. Effect of tryptophan depletion by MGO for 1 day and 7 days.

| Sample | Concentration (µM)  Of MGO  ; 1 day | Concentration (µM)  of MGO  ; 7 days |
| --- | --- | --- |
| Control | 0 | 0 |
| Normal; MGO 1 mM | 1000.00 ± 47.62^###^ | 1000.00 ± 61.65^###^ |
| MGO 1 mM + Tryptophan 1 mM (1 : 1) | 239.51 ± 13.38^***^ | 7.10 ± 0.96^***^ |

All data are shown as means ± SEM. N = 3 (##p < 0.01, ###p < 0.001 vs. Control, ***p < 0.001 vs. Normal).

Table S4. Up- or down-regulated genes in MGO 500 µM treated group.

The table indicates gene expression fold change obtained by microarrays.

| Symbol | Gene name | Log_2_ (fold change) |
| --- | --- | --- |
| **Long-term depression** |  |  |
| Crhr1 | Corticotropin-releasing hormone receptor 1 | 2.46 |
| Gria2 | Glutamate receptor, ionotropic, AMPA2 (alpha 2) | -2.27 |
| Itpr1 | Inositol 1,4,5-trisphosphate receptor 1 | 2.33 |
| Ryr1 | Ryanodine receptor 1, skeletal muscle | -2.32 |
| **Cholinergic synapse** |  |  |
| Chrnb4 | Cholinergic receptor, nicotinic, beta polypeptide 4 | -2.93 |
| Gng8 | Guanine nucleotide-binding protein (G protein), gamma 8 | -2.89 |
| **Purinergic receptor** |  |  |
| P2rx3 | Purinergic receptor P2X, ligand-gated ion channel, 3 | -2.75 |
| **Inflammatory** |  |  |
| Traf6 | TNF receptor-associated factor 6 | 2.40 |
| **tRNA synthetases** |  |  |
| Aars | Alanyl-tRNA synthetase | -2.01 |
| **Amino acid metabolism** |  |  |
| Fbxl22 | F-box and leucine-rich repeat protein 22 | -2.18 |
| Fhit | Fragile histidine triad gene | -3.07 |
| Gfpt2 | Glutamine fructose-6-phosphate transaminase 2 | -2.27 |
| Hrct1 | Histidine rich carboxyl terminus 1 | 2.53 |
| Lrrc43 | Leucine rich repeat containing 43 | -2.71 |
| Lrrc55 | Leucine rich repeat containing 55 | -2.04 |
| Lrrn2 | Leucine rich repeat protein 2, neuronal | 2.02 |
| Padi2 | Peptidyl arginine deiminase, type II | -2.31 |
| Pelp1 | Proline, glutamic acid and leucine rich protein 1 | 2.87 |
| Prrt3 | Proline-rich transmembrane protein 3 | 2.35 |
| Ptdss2 | Phosphatidylserine synthase 2 | 2.40 |
| Ptprt | Protein tyrosine phosphatase, receptor type, T | -8.83 |
| Sdsl | Serine dehydratase-like | 2.22 |
| Sepsecs | Sep (O-phosphoserine) tRNA:Sec (selenocysteine) tRNA synthase | 2.27 |
| Serpinb1a | Serine (or cysteine) peptidase inhibitor, clade B, member 1a | 4.69 |
| Serpinh1 | Serine (or cysteine) peptidase inhibitor, clade H, member 1 | -2.32 |
| Styxl1 | Serine/threonine/tyrosine interacting-like 1 | -2.22 |
| Tha1 | Threonine aldolase 1 | 2.22 |
| Thg1l | tRNA-histidine guanylyltransferase 1-like (S. cerevisiae) | 2.05 |
| Trdmt1 | tRNA aspartic acid methyltransferase 1 | 3.35 |
| Ttll11 | Tubulin tyrosine ligase-like family, member 11 | -2.45 |
| Tyrobp | TYRO protein tyrosine kinase binding protein | -7.20 |
| **Rho family** |  |  |
| Gadd45a | Growth arrest and DNA-damage-inducible 45 alpha | -2.17 |
| Gas1 | Growth arrest specific 1 | 2.07 |
| Gdf9 | Growth differentiation factor 9 | -2.21 |
| Mst1 | Macrophage stimulating 1 (hepatocyte growth factor-like) | -2.53 |
| Srgap1 | SLIT-ROBO Rho GTPase activating protein 1 | -2.17 |
| **Cytoskeleton** |  |  |
| Acta1 | Actin, alpha 1, skeletal muscle | -2.12 |
| Acta2 | Actin, alpha 2, smooth muscle, aorta | -2.00 |
| Coro1a | Coronin, actin binding protein 1A | 3.02 |
| Fgfr1 | Fibroblast growth factor receptor 1 | 2.11 |
| Lrp1 | Low density lipoprotein receptor-related protein 1 | -2.00 |
| Mical2 | Microtubule associated monooxygenase, calponin and LIM domain containing 2 | 3.51 |
| Mical3 | Microtubule associated monooxygenase, calponin and LIM domain containing 3 | 2.18 |

* Values are the means of two independent arrays
